# *De-novo* inter-regional coactivations of pre-configured local ensembles support memory

**DOI:** 10.1101/2021.02.03.429684

**Authors:** Hiroyuki Miyawaki, Kenji Mizuseki

**Affiliations:** Department of Physiology, Osaka City University Graduate School of Medicine, Asahimachi 1-4-3, Abeno-ku, Osaka, 545-8585, JAPAN

**Keywords:** Memory consolidation, Memory acquisition, Memory retrieval, Fear conditioning, Basolateral amygdala, Ventral hippocampus, Pre-limbic cortex, non-REM sleep, Sharp-wave ripples, Cell assembly reactivation

## Abstract

Neuronal ensembles in the amygdala, ventral hippocampus, and prefrontal cortex are involved in fear memory; however, how the inter-regional ensemble interactions support memory remains elusive. Using multi-regional large-scale electrophysiology in the afore-mentioned structures of fear-conditioned rats, we demonstrated that local ensembles activated during fear memory acquisition were inter-regionally coactivated during subsequent sleep, which relied on brief bouts of fast network oscillations. During memory retrieval, coactivations reappeared, accompanying fast oscillations. Ensembles contributing to inter-regional coactivation were configured prior to memory acquisition in the amygdala and prefrontal cortex but developed through experience in the hippocampus. Our observation suggests that elements of a given memory are instantly encoded within various brain regions in a pre-configured manner, whereas hippocampal ensembles and the network for inter-regional integration of the distributed information develop in an experience-dependent manner to form a new memory, which is consistent with hippocampal memory index hypothesis.

## Introduction

Animals acquire memory through experience during wakefulness, and in the subsequent sleep, the acquired labile memory is transformed into a stable form by a process called memory consolidation (Buzsáki, 1989; Klinzing et al., 2019). Cell ensembles within local circuits activated at the time of memory acquisition become also active at the time of memory retrieval (Tonegawa et al., 2015), implying that memory-encoding cell ensembles are maintained stably through memory consolidation. In contrast, memory-responsible brain regions shift with time (Kim and Fanselow, 1992; Squire, 1986), suggesting that the global circuit dynamically changes during memory consolidation. To date, it remains unknown how memory encoding ensembles are inter-regionally interact during memory consolidation and retrieval. Moreover, neuronal activity patterns during awake periods are spontaneously reactivated during the subsequent non-rapid eye movement sleep (NREM) epochs in various brain regions (Girardeau et al., 2017; Peyrache et al., 2009; Wilson and McNaughton, 1994), and this reactivation has an essential role in memory consolidation (Buzsaki, 2015; Ego-Stengel and Wilson, 2010; Girardeau et al., 2009; Maingret et al., 2016). However, it remains controversial whether sleep reactivation occurs synchronously (Girardeau et al., 2017; Ji and Wilson, 2007; Lansink et al., 2009; Olafsdottir et al., 2016; Peyrache et al., 2009; Qin et al., 1997) or independently (O’Neill et al., 2017) in different brain regions.

Ensemble reactivations in the dorsal hippocampus (dHPC) occur during short bouts of fast (100–250 Hz) oscillations in hippocampal local field potentials (LFPs), known as sharp-wave ripples (SWRs) (Buzsaki, 2015). Co-occurrence of SWRs and cortical oscillatory events, such as sleep spindles (9–18 Hz) or cortical ripples (cRipples; 90–180 Hz), has been proposed as the inter-regional information transfer mechanism (Khodagholy et al., 2017; Maingret et al., 2016). SWRs are also observed in the ventral hippocampus (vHPC), while SWRs in dHPC and vHPC occur largely asynchronously, have distinct physiological properties (Patel et al., 2013), and affect activity in downstream regions differently (Sosa et al., 2020). Moreover, fast (90–180 Hz) oscillations are also observed in the amygdala where the oscillatory events are referred to as highfrequency oscillations (HFOs) (Ponomarenko et al., 2003). However, how fast network oscillations in various regions control ensemble reactivations and whether these oscillations regulate interregional communication remain elusive (Skelin et al., 2019). In addition, it has also been proposed that cortical delta (0.5–4 Hz) waves facilitate information transfer by coordinating occurrence timings of various oscillatory events (Klinzing et al., 2019), but the supporting evidence is largely from studies on information transfer between dHPC and neocortical regions. Hence, it remains unknown whether fast oscillations in brain regions other than dHPC are coordinated by delta waves and whether delta waves support inter-regional ensemble communications.

In addition to ensemble reactivation following experience, recent studies have indicated that ensemble activity similar to that during behavior exists prior to experience (Dragoi and Tonegawa, 2011; Farooq et al., 2019; Grosmark and Buzsaki, 2016) (but see (Silva et al., 2015)). Furthermore, whether inter-regional ensemble coordination also exist prior to experience, and cells contributing to the coordination are intrinsically distinct remain to be determined.

In this study, we aimed to investigate inter-regional interactions of local ensemble activities and sought their regulation mechanisms and physiological functions in memory process using fear conditioning as a model. Fear memory involves the vHPC CA1 region (vCA1), basolateral amygdala (BLA), and prelimbic cortex (PL) (Tovote et al., 2015). These brain regions are anatomically interconnected, but a direct projection from the PL to the vCA1 is lacking (Tovote et al., 2015). By simultaneous recordings in the vCA1, BLA, and PL, we suggest that elements of a fear memory are instantly encoded in pre-configured local ensembles, and *de novo* interregional ensemble coactivations bind these elements together and support memory retrieval.

## Results

### Simultaneous recording of neuronal activity from multiple single cells in vCA1, BLA, and PL of fear-conditioned rats

We performed simultaneous large-scale electrophysiological recordings in the vCA1, BLA, PL layer 5 (PL5), and adjacent regions (central amygdaloid nucleus, lateral amygdala, pyriform cortex, bed nucleus of the stria terminalis intra-amygdaloid division, vHPC CA3 region, and ventral subiculum; Figures 1A, 1B and S1) and examined LFPs and 1,220 well-isolated units (Table S1) in 15 freely moving rats. Recordings were performed continuously throughout baseline, conditioning, context retention, cue retention and extinction, retention of extinction, and homecage sessions preceding, interleaved, and following the behavioral sessions (Figures 1C and 1D). The proportion of time spent in freezing behavior indicated that the rats had learned an association between cues and shocks, and they retrieved the association during retention sessions (Figure 1C).

**Figure 1.**
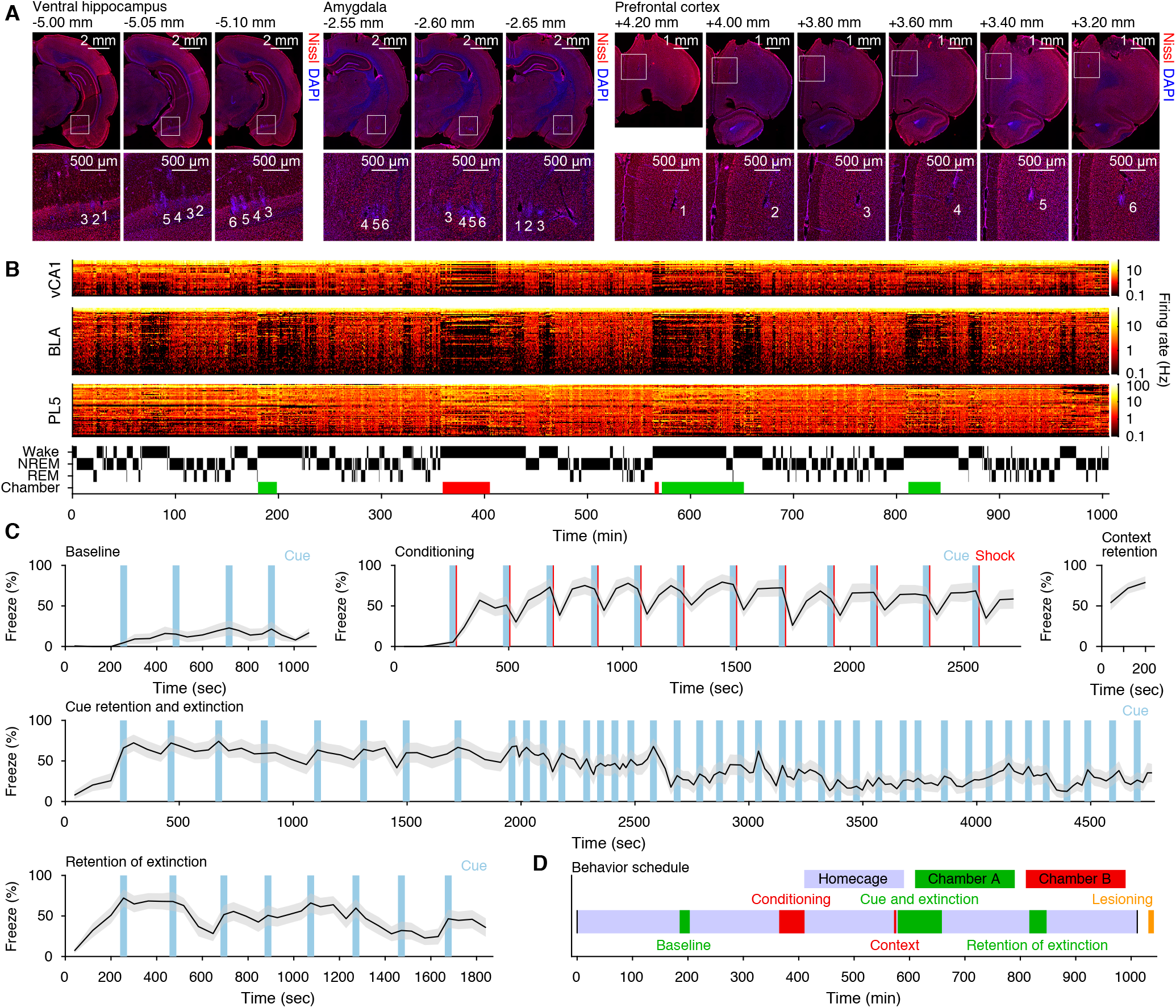
Multi-regional large-scale electrophysiological recording on fear-conditioned rats. **(A)** Locations of electrode tips in a representative example rat. Areas indicated with white squares on the top panels are shown on the bottom in higher magnification. Numbers indicate shank indices within each probe. AP-axis coordinate from the bregma is shown at the top-left of each micrograph. **(B)** Unit activity and hypnogram from the same rat. Each row represents the firing rates of individual units in 10-s bins. Periods of behavioral sessions in safe and shock chambers are indicated with green and red bands on the bottom, respectively. **(C)** The fraction of time in freezing in each behavioral session. Black lines and grey sheds indicate mean and standard error (SE), respectively (n = 15 rats). **(D)** Time schedule of the experiments. Black vertical lines indicate start and end time of recordings. See also Figure S1.

### Memory encoding ensembles in different brain regions are synchronously reactivated during NREM sleep after fear conditioning

First, to investigate whether reactivations of memory encoding ensembles in different brain regions interact, we identified neuronal ensembles, which reflect prominent cofiring of multiple neurons within a short time window (20 ms), in each brain region using independent component analyses (ICA) on spike trains during conditioning sessions (Giri et al., 2019; Lopes-dos-Santos et al., 2013). Then, we estimated instantaneous ensemble activation strength both in pre-conditioning (pre-cond.) and post-conditioning (post-cond.) homecage sessions, which reflects how cofiring patterns within each time bin are similar to those observed during behavior (Giri et al., 2019; Lopes-dos-Santos et al., 2013; Peyrache et al., 2009) (Figures 2A and 2B, Table S2). Inter-regional interactions of ensembles were assessed with cross-correlogram (CCG) analyses of instantaneous ensemble activation strength (Figures 2C, 2D, and 3A). The interregional synchronous ensemble activation during NREM was significantly enhanced after fear-conditioning in BLA–PL5 and vCA1– PL5 pairs (Figures 2C–2E, 3A–3C, Tables S3). These coactivations were detected in individual rats if enough number of ensemble pairs were examined (Table S4). Although there were few negatively correlated pairs (Figures 2D, 2E, 3A, and 3B, Tables S3), herein, we focused on positively correlated pairs. Hereafter, we refer to ensemble pairs showing significant coactivation during post-cond. NREM as coupled ensemble pairs. A few vCA1–BLA coupled ensemble pairs (4 out of 257 pairs) were also identified, but the change was not significant at the population level (Figures 3B and 3C). Similarly, we identified a few coupled ensemble pairs in other region pairs (Figure S2A, Table S3), but the fraction changed from preto post-cond. NREM was not significant. In contrast to the findings in NREM, fractions of coactivated ensemble pairs did not differ between pre- and post-cond. rapid eye movement (REM) sleep (Figures S2B–S2D).

**Figure 2.**
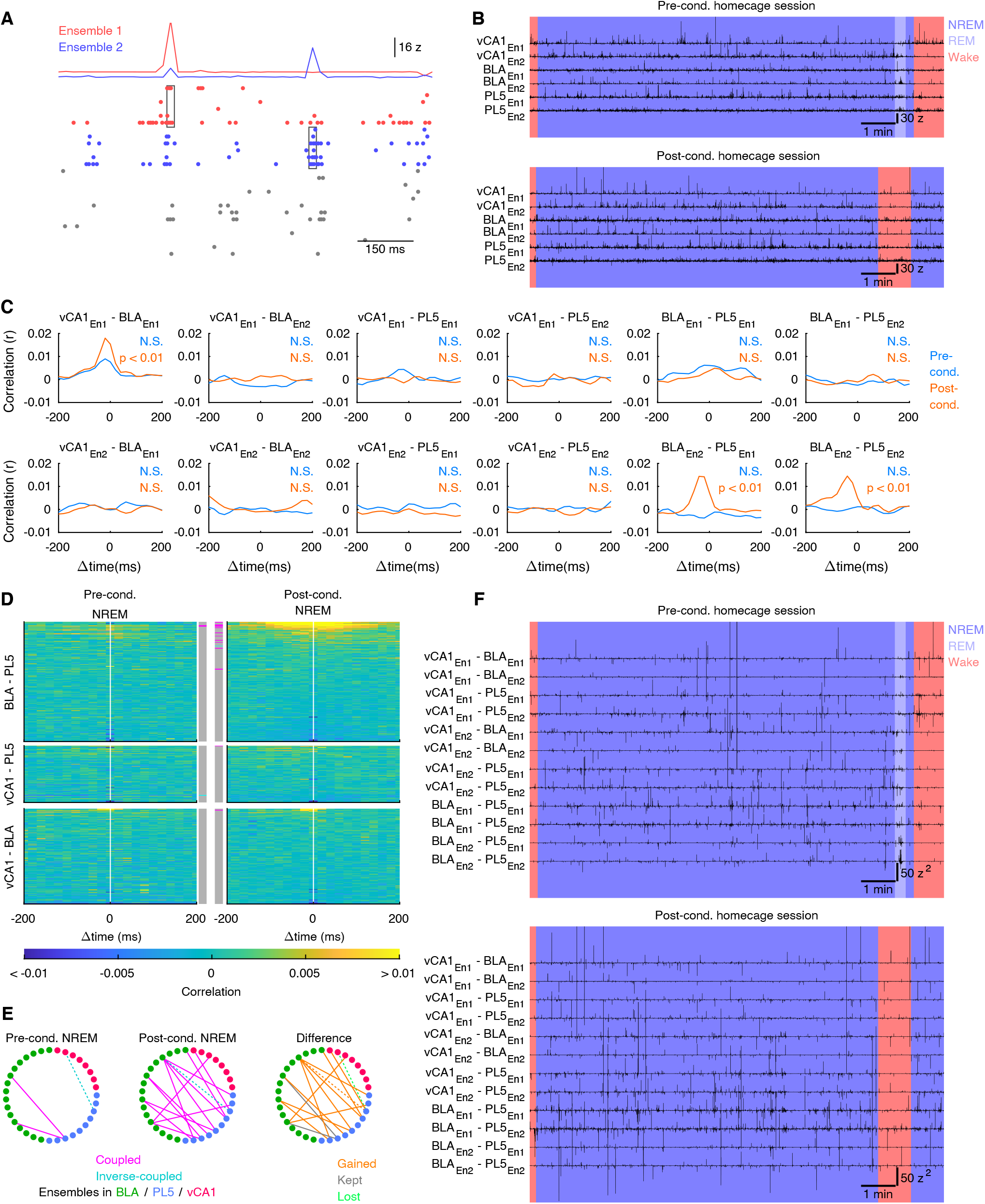
Representative examples of ensemble reactivations and inter-regional coactivations. **(A)** A representative example of vCA1 spike raster plot from the rat presented in Figures 1A and 1B. Two examples of instantaneous ensemble activation strength traces detected from the spike trains are shown on top. Ensembles (group of cofiring neurons in the same region) were identified in the conditioning session, and cells with major contribution to the ensembles (top 6 neurons per ensemble) are highlighted with colors, and spikes of these cells at significant ensemble activation events are highlighted with boxes. **(B)** Instantaneous activation strength of local ensembles in pre- and post-cond. homecage sessions detected from the same rat. Two representative ensembles from each region are shown. Background colors indicate behavioral states. **(C)** Inter-regional CCGs of instantaneous activation strength trances shown in (B) during pre- and post-cond. NREM. Significance of peak on each CCG (determined with shuffling) is superimposed on top-right. **(D)** All inter-regional CCGs of instantaneous activation strength of ensembles obtained from the same rat (n = 170, 80, and 136 for BLA–PL5, vCA1–PL5, and vCA1–BLA ensemble pairs, respectively). Each row represents the CCG for one inter-reginal ensemble pair. Ensemble pairs are sorted based on peak heights of CCGs during post-cond. NREM. Colored bars on middle indicate pairs with significant peaks (magenta) or troughs (cyan) determined with shuffling (p < 0.01). **(E)** Diagrams showing coactivation networks among ensembles obtained from the same rat. Solid and dashed lines indicate ensemble pairs with significant CCG peaks (coupled ensemble pairs) or trough (inverse-coupled ensemble pairs), respectively. The right panel illustrates changes from pre- to post-cond. NREM. **(F)** Instantaneous coactivation strength of inter-reginal ensemble pairs shown in (B).

**Figure 3.**
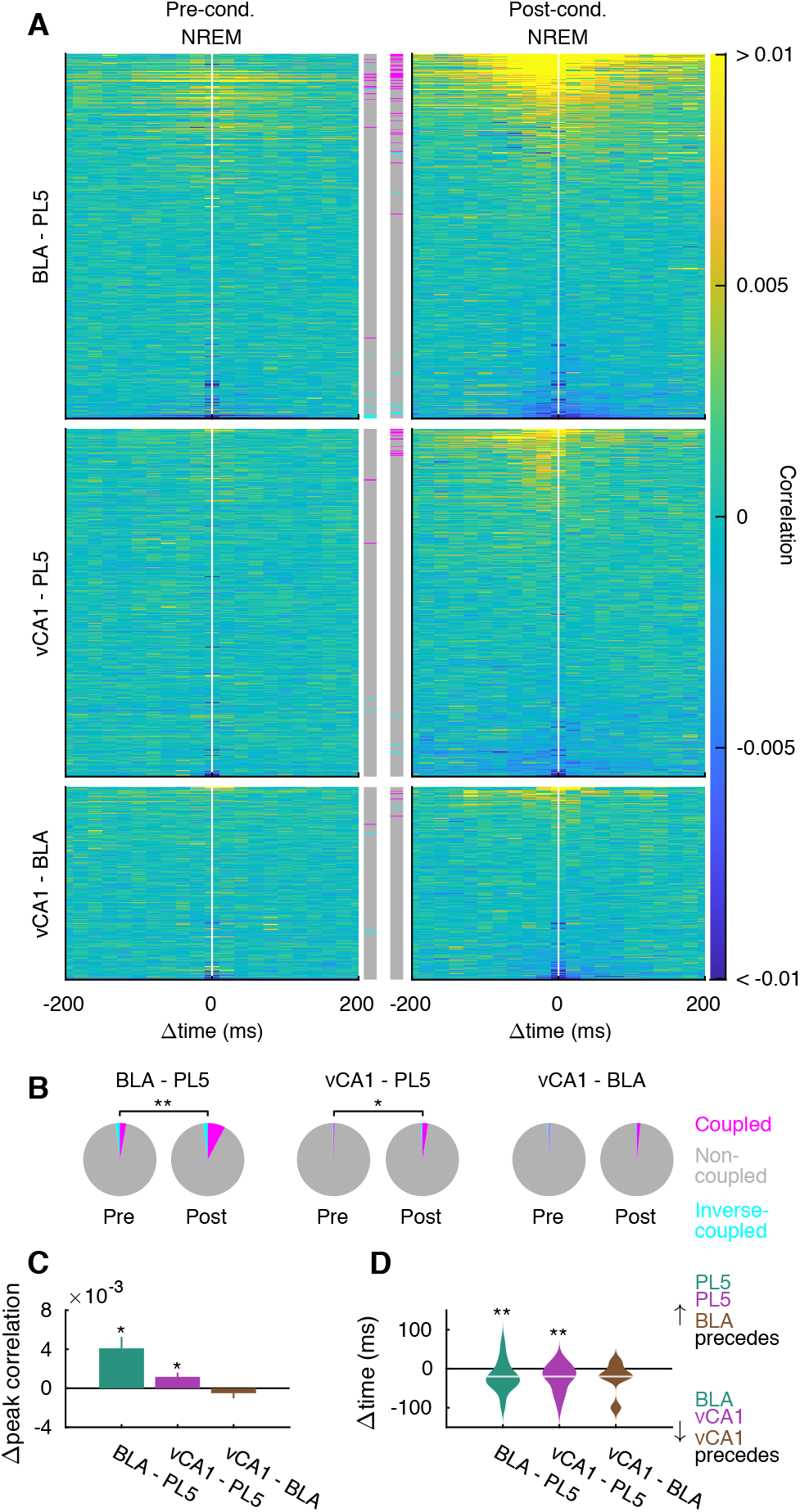
Memory encoding ensembles in different brain regions are synchronously reactivated during non-REM sleep after fear conditioning. **(A)** Inter-regional CCGs of instantaneous activation strength of local ensembles as in Figure 2B, but for data pooled across rats. **(B)** Fractions of ensemble pairs with significant peaks/troughs shown in (A). ** p < 0.01, * p < 0.05, χ^2^ test. **(C)** Changes in CCG peak heights from pre- to post-cond. NREM for all ensemble pairs shown in (A). Error bars indicate SE. * p < 0.05, WSR-test. **(D)** Violin plots of peak time on CCGs with significant peaks. Horizontal bars indicate the median. ** p < 0.01, WSR-test. The numbers of tested pairs and those in each rat are summarized in tables S3 and S4. See also Figure S2.

Distribution of CCG peak time among coupled ensemble pairs showed that reactivation in PL5 ensembles tended to follow that in BLA (20.4 ± 5.7 ms, n= 38 pairs) and vCA1 (22.9 ± 7.4 ms, n = 12 pairs) (Figure 3D). These temporal delays are congruent with monosynaptic transmission latency in long-range projections, such as projections to the prefrontal cortex from vHPC (Degenetais et al., 2003) and BLA (Pérez-Jaranay and Vives, 1991), suggesting that direct projections from vCA1/BLA to PL5 support the identified inter-regional coactivations.

Since emotionally arousing experiences are remembered better than neutral ones (McGaugh, 2004; Paz and Pare, 2013), we hypothesized that a strong aversive experience enhances the inter-regional ensembles coactivation. Thus, to determine whether the coactivation of ensemble pairs also occurs after the baseline session, in which rats are exposed to novel environments and tones without electrical shocks, we identified neuronal ensembles in the baseline session. The fraction of coactivated ensemble pairs did not change between pre- and post-baseline NREM sessions (Figures S2E–S2G). These results indicate that a subset of neuronal ensembles becomes coactivated across brain regions selectively after fear conditioning.

### Amygdalar HFOs, hippocampal SWRs, and prelimbic cRipples contribute to inter-regional ensemble coactivation during NREM

Next, we sought network activity patterns during which the ensemble coactivation among vCA1, BLA, and PL5 occurred. Visual inspection suggested that BLA–PL5 coactivation accompanied fast (∼130 Hz) oscillations in BLA LFP (Figure 4A), known as amygdalar HFOs (Ponomarenko et al., 2003) (Figure S3). HFOs were partially coupled with SWRs and cRipples (14.3 ± 2.5% and 6.0 ± 1.3% of HFO peaks were detected within ± 100 ms periods of SWR and cRipple peaks, respectively; Figures S3C and S3D). HFOs strongly modulated cell firing in the amygdala and other regions (Figure S3E) and enhanced ensemble activations in the BLA (Figure S3F). Similarly, vCA1 and PL5 ensemble activation strength transiently increased at SWR- and cRipple-peaks, respectively (Figure S3F).

**Figure 4.**
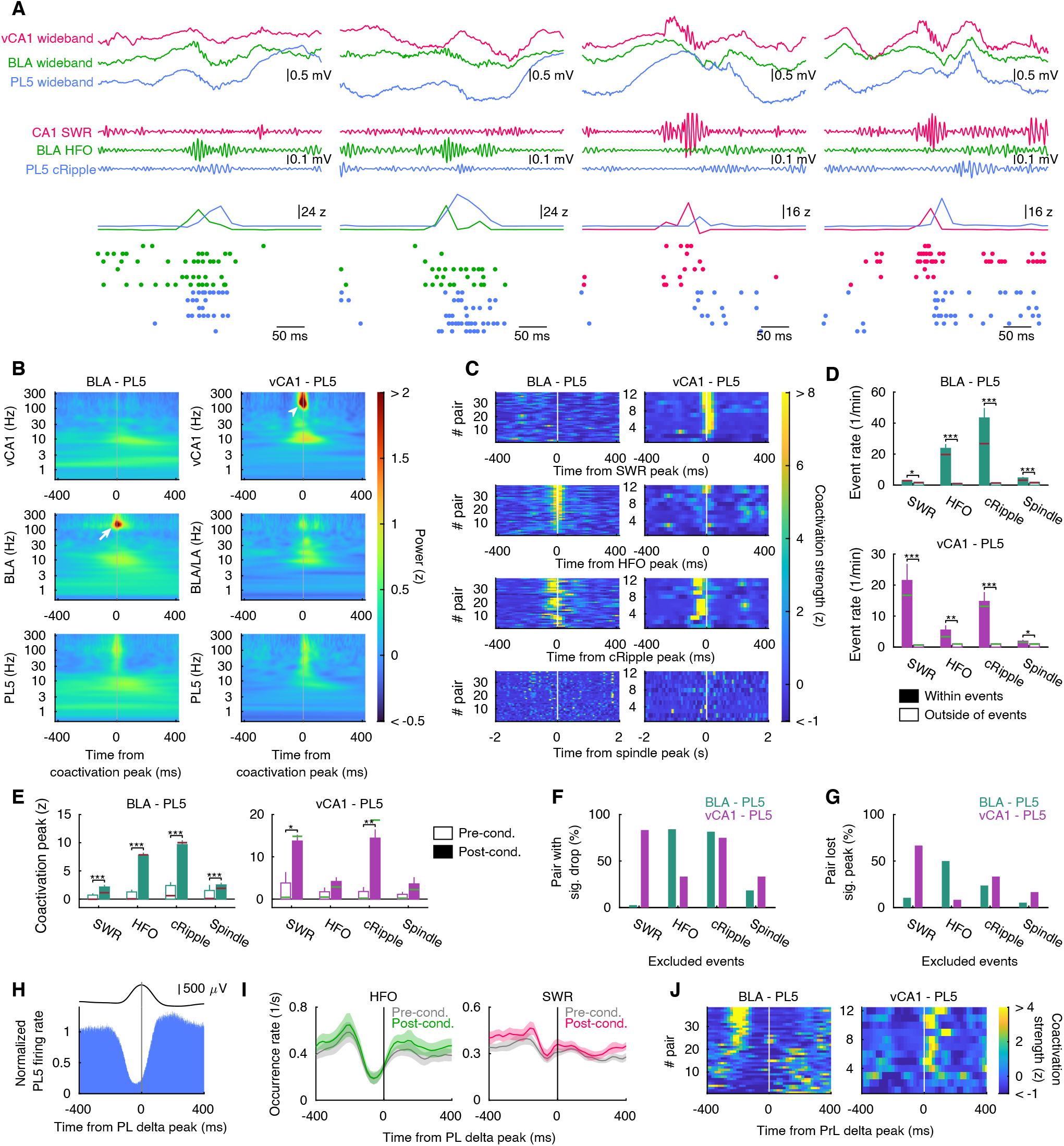
HFOs, SWRs, and cRipples are crucial for inter-regional ensemble coactivations during NREM. **(A)** Representative examples of BLA–PL coactivation (left 2 panels) and vCA1–PL5 coactivation (right 2 panels). Instantaneous ensemble activation strength and spikes from cells with major contributions (top 6 neurons per ensemble) are shown on bottom. **(B)** BLA–PL5 and vCA1–PL5 coactivation-triggered average of LFP wavelet powers during post-cond. NREM. Peaks corresponding to HFOs (white arrow) and SWRs (white arrowhead) are observed at BLA–PL5 and vCA1–PL5 coactivations, respectively. **(C)** Oscillatory event peak triggered average of BLA–PL5 and vCA1–PL5 coactivation strength. The BLA–PL5 and vCA1–PL5 ensemble pairs are sorted based on peak height around time 0. **(D)** Mean and SE of coactivation event rates within/outside of oscillatory events during post-cond. NREM. *** p < 0.001, ** p <0.01, * p < 0.05, WSR-test. Horizontal bars indicate median. **(E)** Mean and SE of coactivation strength peak within oscillatory events during pre- and post-cond. NREM. *** p < 0.001, ** p <0.01, * p < 0.05, WSR-test. Horizontal bars indicate median. **(F, G)** Fractions of coupled ensemble pairs that significantly (p < 0.01, random jittering analyses) reduced CCG peak height (F) and fractions of coupled ensemble pairs that lost significant peaks on CCG (G) by excluding time bins containing SWRs, HFOs, cRipples, or spindles. **(H)** Delta peak triggered average of normalized population firing rate in PL5 (in 1-ms bins, n = 8 rats). Mean PL5 LFP waveforms are presented on top. **(I)** Delta peak triggered average of HFO and SWR occurrence rates. Lines and shades indicate mean and SE (n = 15 / 14 rats for HFOs and SWRs, respectively). **(J)** Delta peak triggered average of BLA–PL5 and vCA1–PL5 coactivation strength. The numbers of analyzed pairs in B–G and J are summarized in Table S3. See also Figures S3 and S4.

Additionally, vCA1–PL5 ensemble co-activation were frequently coincident with SWRs (Figure 4A), while cRipples also cooccurred frequently with BLA–PL5 and vCA1–PL5 ensemble coactivations (Figure 4A). These observations imply tight coupling between inter-regional ensemble coactivations and fast oscillations such as HFOs, SWRs, and cRipples.

To quantify these observations, we calculated instantaneous co-activation strength (Figure 2F) and detected individual coactivation events and obtained ensemble coactivation triggered average of LFP wavelet power (Figure 4B). We detected strong peaks ∼130 Hz in BLA wavelet power at BLA–PL5 ensemble coactivations, which reflect coincidence of BLA–PL5 ensemble coactivation and HFOs. At vCA1–PL5 ensemble coactivations, strong peaks of ∼150 Hz and broad peaks of ∼15 Hz were observed in hippocampal wavelet power, which correspond to ripples and accompanying sharp waves (Oliva et al., 2018) in vHPC. In addition, we found peaks ∼130 Hz of PL5 LFP at BLA–PL5 and vCA1–PL5 ensemble coactivations (Figure 4B). Consistent with this, coactivation events transiently increased within cRipples (Figures 4C and 4D). Furthermore, BLA– PL5 and vCA1–PL5 ensemble coactivation were transiently enhanced at HFOs and SWRs, respectively (Figures 4C and 4D). These enhancements were more prominent in post-than in pre-cond. NREM (Figure 4E). In contrast, occurrence rates of HFOs and SWRs did not significantly change and those of cRipples increased moderately between pre- and post-cond. NREM (Figure S4E), indicating that development of inter-regional ensemble coactivation (Figures 2 and 3) are due to enhancement of ensembles coactivations during fast oscillations, but not to increase of fast oscillation events. These results suggest that HFOs, SWRs, and cRipples contribute to inter-regional ensemble coactivations.

To clarify the contributions of HFOs, SWRs, cRipples, and spindles in the coactivations, we excluded time bins that contained these oscillatory events and then reperformed CCG analyses of instantaneous ensemble activation strength (Figures S4A–S4D). We found that 84.2% of BLA–PL5 and 83.3% of vCA1–PL5 coupled ensemble pairs displayed significant CCG peak reduction when HFOs and SWRs were excluded, respectively (Figure 4F). Similarly, exclusion of cRipples significantly decreases CCG peaks in 81.6 % of BLA– PL5 pairs and 75.0 % of vCA1–PL5 pairs (Figure 4F). Further, the peaks on CCGs were no longer significant after the exclusion of HFOs/SWRs in 50.0% of BLA–PL5 and 66.7% of vCA1–PL5 coupled ensemble pairs, respectively (Figure 4G). Removal of cRipples resulted in loss of significant peaks in 23.7 % of BLA–PL5 pairs and 33.3 % of vCA1–PL5 pairs. Moreover, ensemble coactivation in vCA1–BLA depended on both SWRs and HFOs (Figure S4F). In contrast, spindle exclusion had only moderate effects (Figures 4F, 4G, and S4F). These results indicate that amygdalar HFOs and hippocampal SWRs contribute to BLA–PL5 and vCA1–PL5 ensemble coactivations during NREM, respectively, and cRipples contribute ensemble coactivation of both region pairs.

### Cortical delta waves coordinate inter-regional ensemble coactivation during NREM

NREM sleep is characterized by slow oscillations (<1 Hz) and delta waves (1–4 Hz), associated with alternation of silent and active periods of large cortical neuronal populations (Steriade et al., 1993a; Steriade et al., 1993b; Steriade et al., 2001). Peaks of delta waves recorded in the deep layers of neocortex are concomitant with generalized silent periods in the neocortex (Figure 4H), known as DOWN states (Luczak et al., 2007; Miyawaki et al., 2017; Steriade et al., 1993b; Vyazovskiy et al., 2009). PL delta also modulated neuronal firing in BLA, but not in vCA1 (Figure S4G). We observed that occurrence rates of HFOs peaked 200 ± 19 ms (n = 15 rats) prior to PL delta peaks in post-cond. NREM (Figure 4I). Consistent with a tight relationship with HFOs (Figures 4A–4G), BLA–PL5 ensemble coactivations were also enhanced at similar timing (241 ± 14 ms prior to delta peaks in post-cond. NREM, n = 38 pairs; Figure 4J). These time gaps were longer than typical duration from delta onsets to peaks (101 ± 3.6 ms in post-cond. NREM, n = 15 rats) but shorter than duration from delta offset to next delta peak (1,008 ± 47 ms), indicating that BLA–PL5 ensemble coactivation is immediately followed by UP–DOWN transitions.

Similar to previous observations in dHPC (Maingret et al., 2016; Molle et al., 2006; Oyanedel et al., 2020; Peyrache et al., 2011; Peyrache et al., 2009; Sirota et al., 2003), vHPC SWR occurrences were moderately increased around UP–DOWN transitions (reached maxima 105 ± 88 ms prior to delta peaks in post-cond. NREM, n= 14 rats; Figure 4I). Surprisingly, vCA1–PL5 coactivation strength of ensembles was enhanced during DOWN states (Figure 4J); coactivation of vCA1–PL5 ensemble pairs were peaked 56 ± 15 ms after delta peaks (n = 12 pairs), where PL was presumably around DOWN-UP transitions (Figure 4H; duration from delta peak to offset was 109 ± 5.4 ms in post-cond. NREM, n = 15 rats). These observations indicate that subsets of SWRs that preferentially occur around DOWN–UP transitions are involved in vCA1–PL5 ensemble coactivations.

In sum, vCA1–PL5 and BLA–PL5 ensemble coactivation preferentially occur at distinct time-lag with respect to delta waves, suggesting that delta waves coordinate timing of ensemble coactivation in a brain-region combination dependent manner.

### BLA–vCA1–PL5 triple-activation enhanced in post-cond. NREM

In addition to coactivation of inter-regional ensemble pairs, we also observed nearly simultaneous activation of BLA, vCA1, and PL5 ensembles (Figure 5A). To quantify this observation, first we expanded CCG analyses for triplets by calculating “triple-CCG” defined as products of three activation strengths with various time shift (Figure 5B). We then tested significance of triple-CCG peaks by chunk shuffling in post-cond. NREM and identified 100 coupled ensemble triplets (out of 2,925 possible triplet combinations). Every single rat with implants in BLA, vCA1, and PL5 had at least one coupled triplet (Table S4). Peak position of triple-CCG varied across triplets but significantly skewed from uniform (p < 0.001, χ^2^ test, n = 100 triplets), and the histogram had a significant peak at [−60 ms, −20 ms] (Figure 5C), suggesting that most commonly vCA1 ensemble activated first then BLA and PL5 ensembles follows in this order.

**Figure 5.**
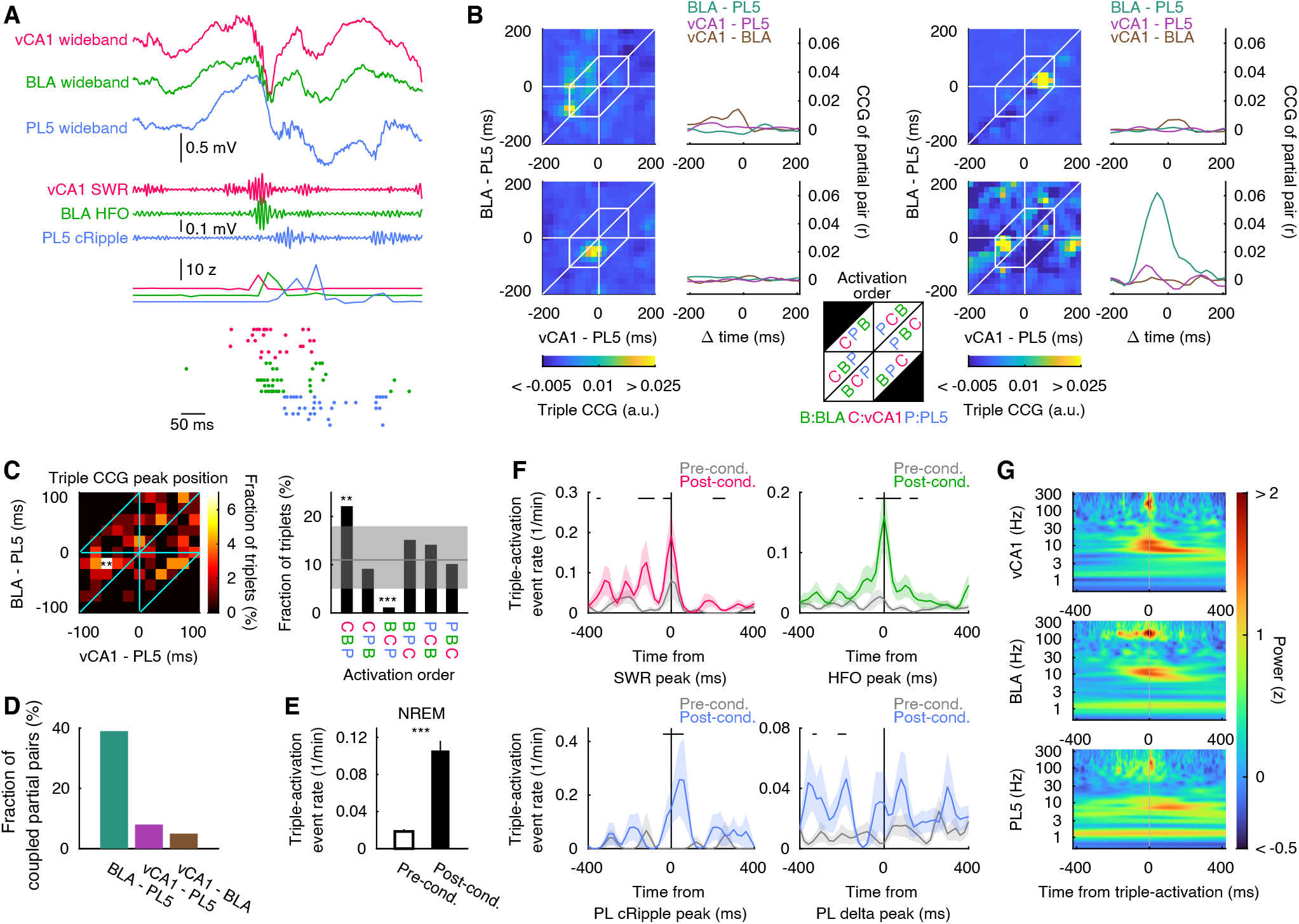
Triple-activation across vCA1, BLA, and PL5 during post-cond. NREM. **(A)** Representative example of triple-activation across vCA1, BLA, and PL5 ensembles. Instantaneous ensemble activation strength and spikes from cells with major contributions (top 6 neurons per ensemble) are shown on bottom. **(B)** Representative examples of triple-CCGs and CCGs of participating ensembles pairs (partial pairs). White hexagons on left panels indicate range for peak detection (time gaps of all combinations < 100 ms). Top left one illustrates the triplet shown in (A). Inset in the middle illustrates order of ensemble activation. **(C)** Distributions of triple-CCG peak position (left) and ensemble activation order (right) across triplets (n = 100 triplets). The peak distribution is non-uniform (p < 0.001, χ^2^ test) with significant peak (** p < 0.01, based on Poisson distribution with Bonferroni correction). Gray line and sheds show mean and 95% confidence interval, respectively, assuming uniform distribution. ** p < 0.01, *** p < 0.001, test based on Poisson distribution. **(D)** Fraction of triplets whose partial pairs had significant peaks on their CCG. **(E)** Mean BLA–vCA1–PL5 triple-activation event occurrence rate per triplets. *** p < 0.001, WSR-test. Error bars indicate SEs (n = 100 triplets). **(F)** Oscillatory event peak triggered average of BLA–vCA1–PL5 triple-activation event rates. Lines and sheds indicate means and SEs (n =100 triplets). Periods with significant differences between pre- and post-cond. are indicated with black ticks on the top (p < 0.05, WSR-test). **(G)** Triple-activation event triggered average of wavelet power (n = 100 triplets) in vCA1, BLA, and PL5.

Next, we examined whether the triple-activations were just coincidence of ensemble coactivation events. We defined partial pairs as pairs of ensembles participating to triplet of interest (each triplet has 3 partial pairs) and found that on average only 17.3 % of partial pairs were coupled (39%, 8.0%, and 5.0% for BLA–PL5, vCA1– PL5, and vCA1–BLA, respectively, n = 100 partial pairs for each; Figures 5B and 5D). This result suggests that triple-activations reflects existence of restricted time windows in which ensembles of the three brain regions are preferentially activated together.

To examine temporal dynamics of triple-activation, we detected triple-activation evens with a method similar to coactivation events detection. First, we calculated instantaneous triple-activation strength as products of instantaneous ensemble activation strength with optimal time shift determined based on triple-CCG peak position, then detected triple-activation events by thresholding the instantaneous triple-activation strength traces. Similar to enhancement of inter-regional coactivation in post-cond. sleep (Figures 2 and 3), triple-activation event rates were significantly increased after fear conditioning (Figure 5E). Enhancement of triple-activation in post-cond. NREM was prominent at SWR-, HFO-, and cRipple-peaks (Figure 5F). Consistently, triple-activation triggered average of LFP wavelet power shows clear peaks corresponding to SWRs, HFOs, and cRipples (Figure 5G). However, delta wave-modulation on triple-activation events were not prominent (Figure 5F), contrasting to those on coactivation of ensemble pairs (Figure 4J). These findings indicate that triple-ensemble activation events were enhanced during post-cond. NREM by SWRs, HFOs, and cRipples.

### BLA–PL5 and vCA1–PL5 ensemble coactivations develop in distinct time courses

We further examined whether the coactivations (Figures 2 and 3) and triple-activations (Figure 5) of ensembles exist prior to the time of memory acquisition or develop after experience. The shock triggered average of coactivation events revealed that coupled ensemble pairs were significantly more coactivated than non-coupled ones in BLA–PL5 at the time of memory acquisition (Figure 6A). Such difference was not detected in vCA1–PL5 ensemble pairs (Figure 6B). Triple-activation of BLA, vCA1, and PL5 ensembles were also enhanced at shock onsets (Figure 6C). BLA ensembles coupled with vCA1 or PL5 were more frequently activated during shock (∼ 2 s) than non-coupled BLA ensembles (Figure S5A), and PL5 ensembles coupled with vCA1 or BLA showed transiently increased activation at shocks onset (Figure S5A). However, the activation strength of vCA1 ensembles was similar between coupled and non-coupled ensembles (Figure S5A). These results indicate that coactivation of BLA–PL5 coupled ensemble pairs and triple-activation of BLA– vCA1–PL5 coupled triplets exist or form rapidly during memory acquisition, but coactivation of vCA1–PL5 coupled ensemble pairs develops in sleep after experiences.

**Figure 6.**
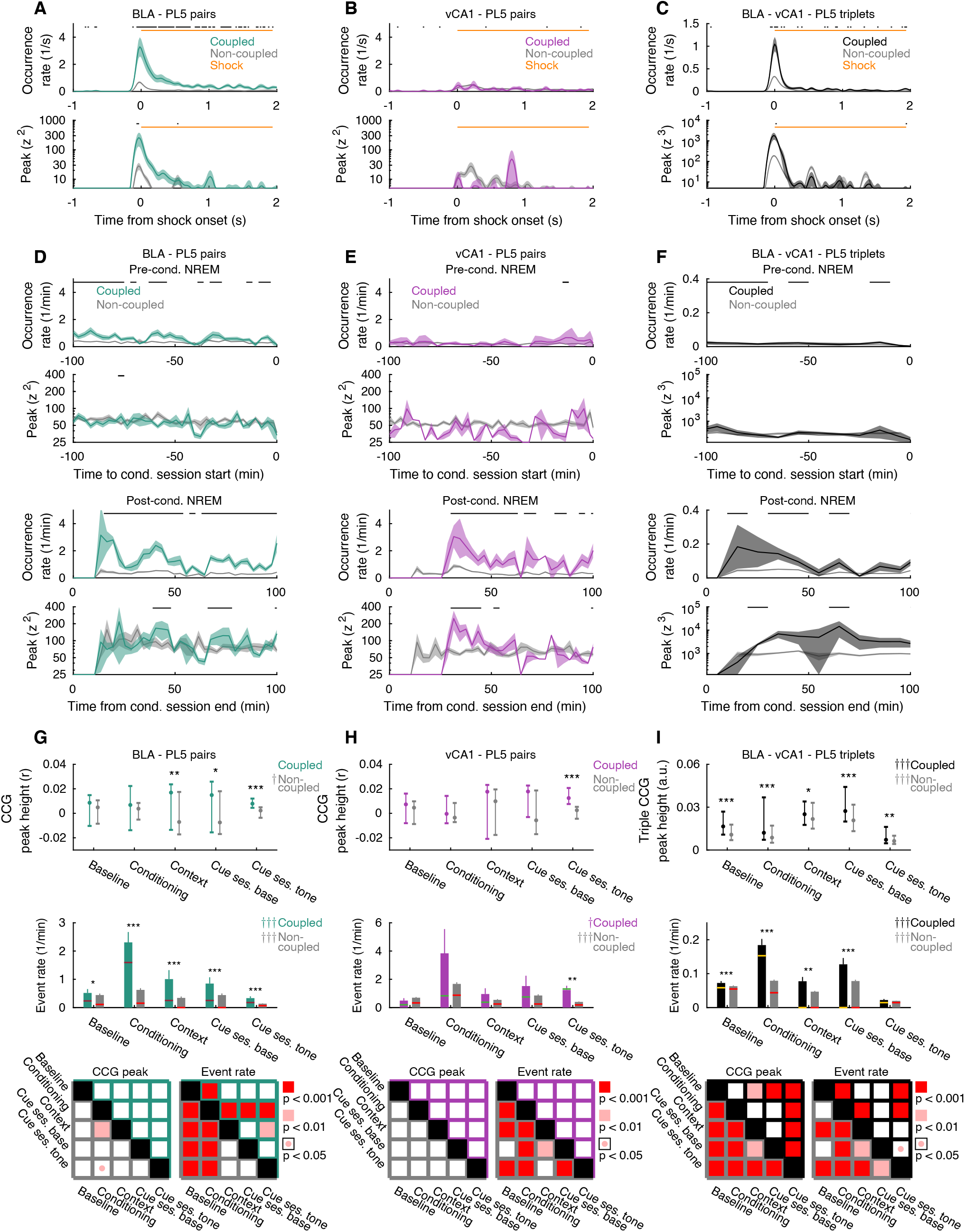
BLA–PL5 and vCA1–PL5 ensemble coactivations develop in distinct time courses. **(A-F)** Shock-triggered average of coactivation/triple-activation events (A-C) and time aligned average of coactivation/triple-activation events during NREM (D-F). Lines and shades represent mean and SE, respectively. Periods with significant differences between coupled and non-coupled ensemble pairs are indicated with black ticks on the top (p < 0.05, WSR-test). **(G-I)** Median of CCG/triple-CCG peak heights and mean of coactivation/triple-activation event rates during behavioral sessions. The cue retention session (Cue ses.) was divided at the onset of the first tone (Cue ses. base and Cue ses. tone, respectively). Top, medians and inter-quadrant ranges (IQR). Middle, mean (bars), SE (error bars), and median (horizontal bars). * p < 0.05, ** p < 0.01, *** p < 0.001, Wilcoxson rank sum test (WRS-test). ††† p < 0.001, †† p <0.01, † p < 0.05, Friedman test. Significance of pair-wise comparison across behavioral sessions are summarized on bottom (Post-hoc WSR-test with Bonferroni correction following Friedman test). Top and bottom halves represent coupled and non-coupled pairs/triplets, respectively. Coupled and non-coupled pairs/triplets separated based on coactivation/triple-activation in post-cond. NREM. The numbers of analyzed pairs and triplets are summarized in Table S3 and S4. See also Figures S5 and S6.

Coactivation events were aligned with the start or end time of conditioning sessions to understand the time evolution of coactivation during NREM (Figures 6D, 6E, and S5B). The occurrence rates of the coactivations of BLA–PL5 coupled ensemble pairs significantly increased from pre- to post-cond. NREM (Figure 6D; Δmean occurrence rate = 0.78 ± 0.14 min^-1^, p < 0.001, WSR-test). In post-cond. NREM, BLA–PL5 coupled ensemble pairs were coactivated more frequently than non-coupled pairs, while the strength of coactivation events was comparable (Figure 6D). Interestingly, similar patterns were also observed in vCA1–PL5 ensemble pairs (Figure 6E). In BLA, reactivations of coupled ensembles were more frequent than those of non-coupled ensembles (Figure S5C). In contrast, coupled and non-coupled ensembles were comparable in terms of occurrence rate and peak strength in vCA1 and PL5 during post-cond. NREM (Figure S5C), suggesting that enhanced coactivations in vCA1–PL5 coupled ensemble pairs are due to time aligned activations (but not increased activation occurrence rate or strength) of the participating ensembles. We also observed increase of triple-activation event rates in coupled ensemble triplets compared to non-coupled ones during post-cond. NREM (Figure 6F). Contrasting to ensemble pairs, peak heights of triple-activation events were also enhanced during post-cond. NREM (Figure 6F). These observations further support the precise time alignments of ensemble activations across brain regions.

We then investigated whether coactivations and triple-activations persist in other behavioral sessions. In BLA–PL5, coactivations of coupled ensemble pairs appeared at conditioning sessions and reappeared in context- and cue-retention sessions (Figures 6G and S6A). Coactivation in BLA–PL layer 2/3 coupled ensemble pairs demonstrated similar patterns (Figures S6A and S6B). In contrast, vCA1–PL5 coupled ensemble pairs became more coactive than non-coupled ones after the first tone in cue retention sessions, and such difference was not detected in the conditioning sessions (Figures 6H and S6A). Coactivation in vCA1–BLA coupled ensemble pairs in behavioral sessions appeared at conditioning sessions (Figure S6C). Importantly, in the cue retention sessions, vCA1–PL5 ensemble coactivations were not detected before the first tone (Figure 6H), suggesting that the ensemble coactivations are associated with memory recall. Collectively, these results indicate that the time evolution of inter-regional ensemble coactivation depends on the participating regions.

Triple-activation of coupled BLA–vCA1–PL5 ensemble triplets stayed stronger and more frequent than non-coupled ensemble triplets across behavioral sessions (Figure 6I). In addition, triple-activation of coupled ensemble triplets were significantly more frequent during conditioning sessions than baseline sessions and cue sessions after first tone onset (Figure 6I), implying that triple-activation is involved in memory acquisition rather than memory retrieval.

### Fast oscillations coordinate inter-regional ensemble coactivation during memory retrieval

Next, we sought network activity patterns during which the coactivation in cue retention sessions occurred. Since triple-activation events were rare during context and cue retention sessions (98%, 91%, and 75% of coupled triplets had no or one triple-activation events during context retention sessions and cue retention sessions before/after first tone onsets, respectively), we did not further analyze triple-activation related activity patterns during these sessions. Similar to post-cond. NREM (Figures 4A–4C), we observed that BLA–PL5 and vCA1–PL5 ensemble coactivations were accompanied by awake HFOs (aHFOs) and SWRs, respectively (Figure 7A). In addition, fast PL oscillations were also observed at the time of coactivations (Figure 7A).

**Figure 7.**
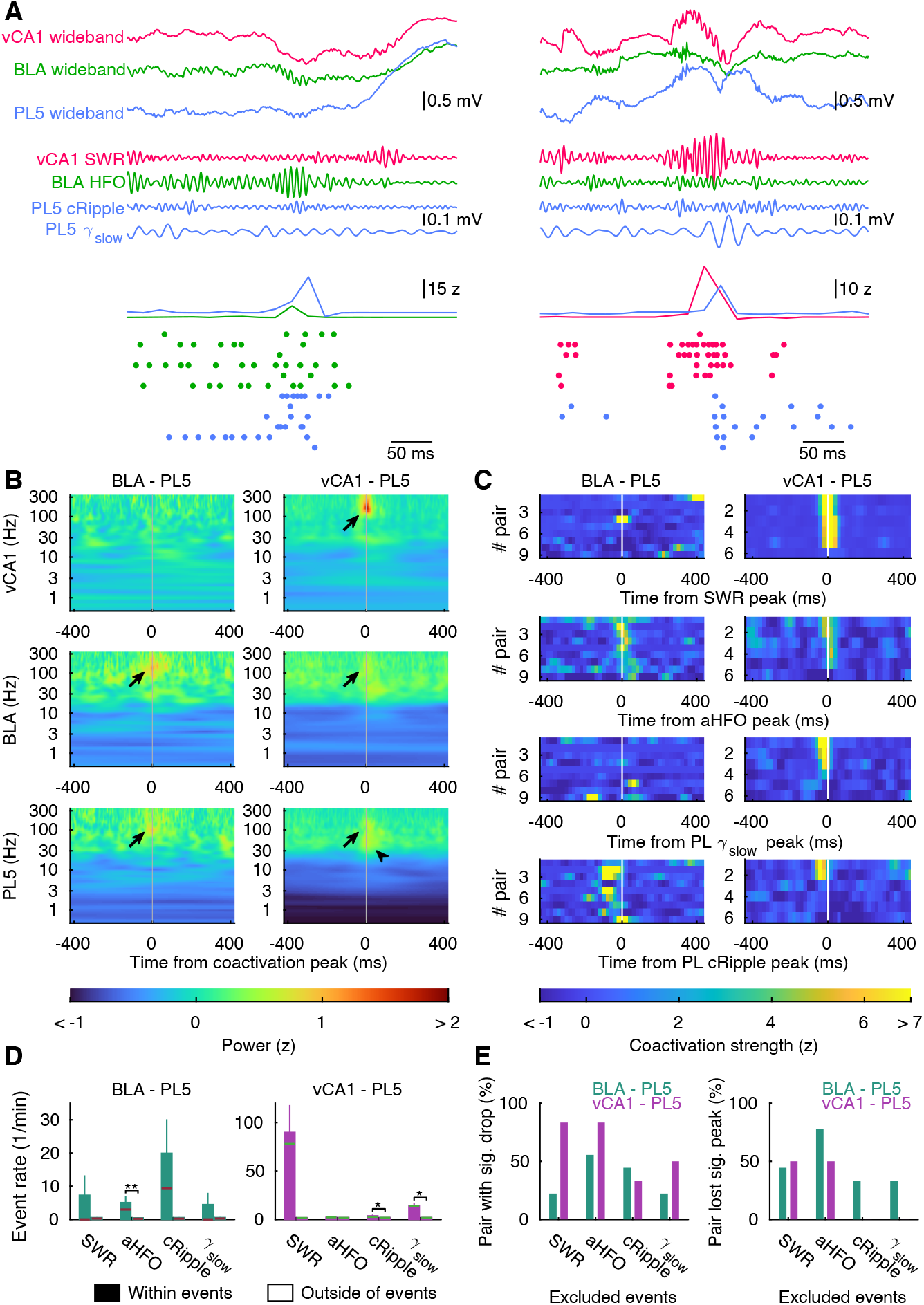
Fast oscillations host coactivation during memory retrieval. **(A)** Representative examples of BLA–PL5 coactivation (left) and vCA1–PL5 coactivation (right) in awake periods. Ensemble activation strength and spikes from cells with large weight are shown on bottom. **(B, C)** Coactivation triggered average of local field potential wavelet power (B) and the oscillatory event-triggered average of instantaneous coactivation strength (C) during cue retention sessions. Only coupled ensemble pairs with significant peaks during cue retention sessions (n = 9 / 6 for BLA–PL5 / vCA1–PL5 ensemble pairs) were used. Black arrows in (B) point peaks reflecting SWRs in vCA1, aHFOs in BLA, and cRipples in PL5, respectively. The black arrowhead on (B) indicates a peak reflecting γslow. **(D)** Mean and SE of coactivation event rates within/outside of oscillatory events during post-cond. NREM. *** p < 0.001, ** p <0.01, * p < 0.05, WSR-test. Horizontal bars indicate median (n = 9 / 6 for BLA–PL5 / vCA1–PL5 ensemble pairs). **(E)** Fractions of coupled ensemble pairs that significantly (p < 0.01, random jittering analyses) reduced CCG peak height and that lost significant peaks on CCG by excluding time bins containing SWRs, aH-FOs, cRipples, or γslow (n = 9 / 6 for BLA–PL5 / vCA1–PL5 ensemble pairs). See also Figure S6.

To better understand the observations, we calculated the ensemble coactivation event-triggered average of LFP wavelet power and detected strong peaks reflecting aHFOs and SWRs at BLA–PL5 co-activation and vCA1–PL5 coactivations, respectively (Figure 7B). Consistently, we observed transient enhancement of BLA–PL5 and vCA1–PL5 ensemble coactivations around aHFO- and SWR-peaks, respectively (Figure 7C). In addition, cRipples tended to follow BLA–PL5 ensemble coactivation (Δt = −35.6 ± 14.5 ms, n = 9 pairs, p = 0.055, WSR-test; Figure 7C). BLA–PL5 coactivation occurrence rates elevated significantly during aHFOs (Figure 7D) and vCA1–PL5 coactivation tended to occur within SWRs (p = 0.063, WSR-test, n = 6 pairs). We also observed that 55.6% of BLA–PL5 coupled ensemble pairs and 83.3% of vCA1–PL5 ensemble pairs dropped their CCG peaks significantly when bins containing SWRs and aHFOs were excluded from the analyses, respectively (Figure 7E). In addition, we observed transient increase of PL5 LFP wavelet power in ∼130 Hz (Figure 7B), which corresponds to cRipples, at both BLA–PL5 and vCA1–PL5 ensemble coactivations. BLA–PL5 and vCA1–PL5 ensemble coactivation occurred more frequently within cRipples (Figure 7D), although difference in BLA–PL5 did not reached statistical significance (p = 0.055, WSR-test, n = 9 pairs). We also observed other PL5 wavelet power peaks in slow gamma (γ_slow_) band (30–60 Hz) at vCA1–PL5 ensemble coactivations (Figure 7B). Consistent with this, peaks of γ_slow_ in PL5 co-occurred with vCA1–PL5 coactivation (Figure 7C) and vCA1–PL5 ensemble coactivation event rates were higher during γ_slow_ epochs (Figure 7D). These findings indicate that inter-regional coactivations during memory retrieval accompany fast oscillations in participated regions. In contrast, no noticeable peaks were detected on PL fast gamma (γ_fast_; 60–90 Hz) triggered average of coactivation strength (Figure S6D). These results suggest that aHFOs and cRipples host BLA–PL5 ensemble coactivations, while awake SWRs and PL5 γ_slow_ host vCA1–PL5 ensemble coactivations during memory retrieval.

### Cell ensembles are configured prior to conditioning in BLA and PL5 but not in vCA1

Recent studies have suggested that memory encoding cells are more excitable prior to experience, and memory encoding ensembles may be configured before experience (Mizuseki and Buzsaki, 2013; Mizuseki and Miyawaki, 2017). To determine whether ensembles and cells contributing to inter-regional coactivation are also pre-determined, we first examined significance of ensemble activation event rates by comparing those of surrogate ensembles (Figure 8A). In BLA and PL5, the observed ensemble activation event rates were significantly higher than chance level in both pre- and post-cond. NREM, indicating that cell ensembles are pre-configured in these regions. In contrast, vCA1 ensembles coupled with PL5 were significantly activated only after fear-conditioning, indicating that co-activation contributing vCA1 ensembles developed in an experience dependent manner.

**Figure 8.**
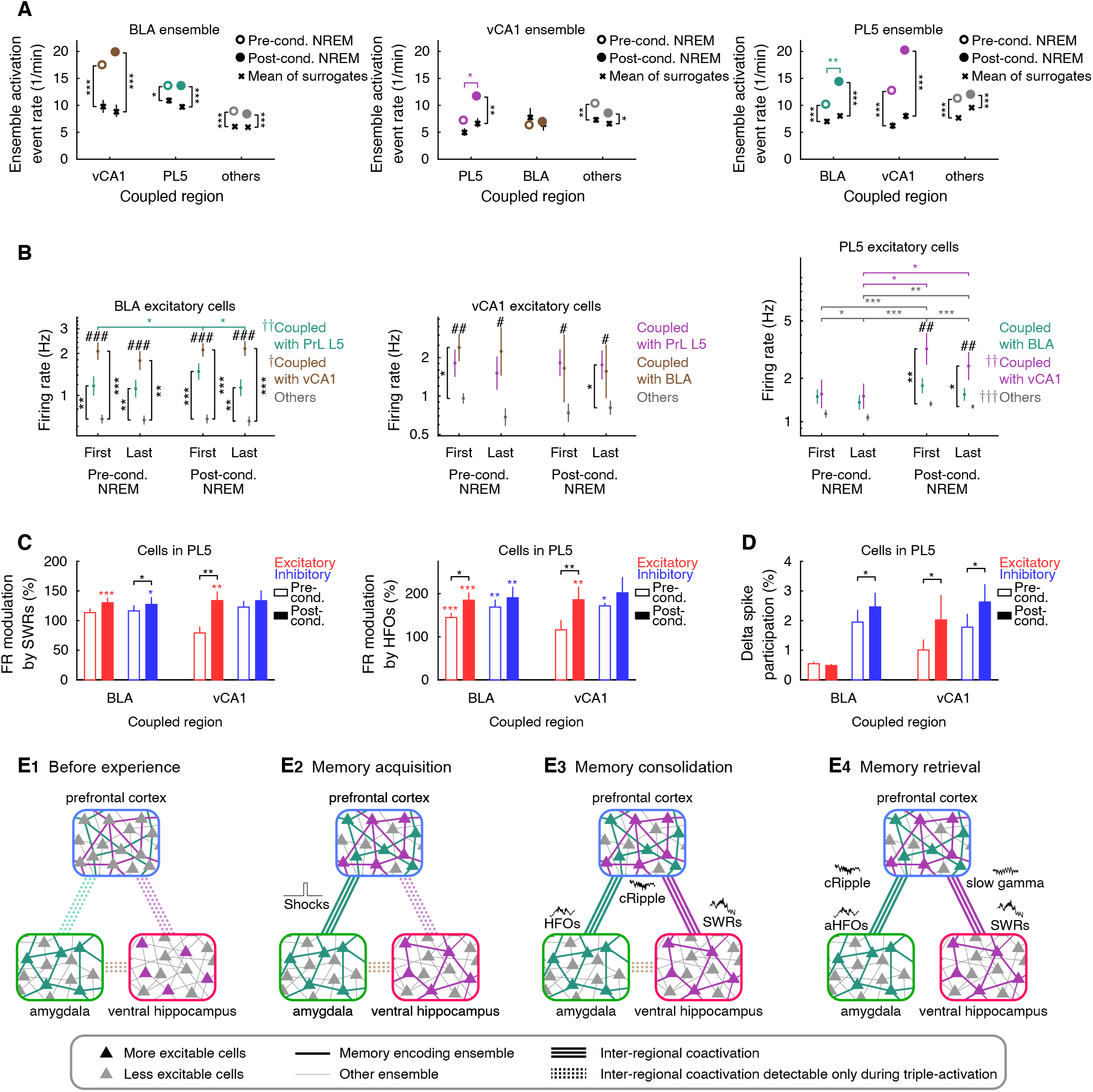
Distinct firing properties of coactivation contributing cells are configured prior to conditioning in BLA and vCA1 but develop after conditioning in PL5. **(A)** Mean ensemble activation event rates in NREM. Ensembles that were not coupled with PL5, BLA, or vCA1 are shown as others. Black crosses and error bars indicate median and IQR of surrogate mean (n = 500 surrogates). * p < 0.05, ** p < 0.01, *** p < 0.001, random shuffling/WSR-test for black/colored marks. **(B)** Mean and SE of firing rates during NREM in homecage sessions. Cells that were not coupled with PL5, BLA, or vCA1 are shown as others. # p < 0.05, ## p < 0.01, ### p < 0.001, Kruskal–Wallis test; † p < 0.05, †† p < 0.01, ††† p < 0.001, Friedman test; * p < 0.05, ** p < 0.01, *** p < 0.001, post-hoc WRS-test/WSR-test with Bonferroni correction for black/colored marks. **(C, D)** Modulation of firing rate by SWRs and HFOs (C) and participation rate of delta spikes (D) for BLA/vCA1–coupled PL5 cells. * p < 0.05, ** p < 0.01, *** p < 0.001 WSR-test for modulation within pre- or post-cond. NREM (C, colored) and changes from pre- to post-cond. NREM (C and D, black). Error bars indicate SE. **(E)** Schematic summary of inter-regional ensemble coactivation development, suggesting that elements of memories are instantly embedded in pre-configured amygdalar and prefrontal cortical ensembles, whereas inter-regional network to bind the distributed information develops through memory consolidation during the following sleep period and reactivated during memory retrieval. The numbers of analyzed ensembles in (A) and cells in (B-D) are summarized in Table S5 and S6, respectively. See also Figures S7 and S8.

Next, we examined whether excitability of cells contributing to the ensemble coactivation are also pre-determined. We defined ensemble coactivation contributing cells from each coupled ensemble as cells with high absolute weight on ICA projection (> 0.3) and defined their coupled region as the partner region of the coupled ensemble. In principle, a given cell can have multiple coupled regions, but most cells had one or no coupled region, except for BLA inhibitory cells (Table S6, Figure S7A). In BLA and vCA1, coactivation contributing excitatory cells, but not inhibitory cells, tended to fire faster than others across brain states (Figure S7B and S7C). The higher firing rates of coactivation contributing excitatory cells in vCA1 and BLA were observed even in NREM during the earlier halves of pre-cond. homecage sessions (Figure 8B). Among ensemble coactivation contributing excitatory cells in BLA and vCA1, firing modulation by fast LFP oscillations was largely unchanged between pre- and post-cond. NREM; the only change detected was HFO modulation of firing in BLA (Figures S7D and S7E). These results suggest that ensemble coactivation contributing excitatory cells are more excitable in BLA and vCA1.

Conversely, ensemble coactivation contributing cells in PL5 developed distinct firing properties after conditioning. Firing rates of PL5 excitatory cells coupled with vCA1 were higher than those of others in post-cond. NREM, whereas firing rates were comparable across PL5 excitatory cells coupled with vCA1/BLA and other cells in pre-cond. NREM (Figure 8B). We also found that SWR/HFO-modulation of ensemble coactivation contributing PL5 excitatory cell firing was enhanced by conditioning (Figure 8C), although changes in SWR modulation in excitatory cells coupled with BLA was not statistically significant (p = 0.069, WSR-test). These results indicate that ensemble coactivation contributing cells in PL5 refine their firing activity through experience, and oscillatory event modulation of their firing is changed by conditioning. We observed that a minority of spikes of PL cells occurred even during delta peaks (Figure 4H), known as delta spikes (Todorova and Zugaro, 2019). Furthermore, PL5 cells coupled with vCA1 participated in delta spikes more frequently in post-cond. NREM than in pre-cond. NREM (Figure 8D); this finding is in line with the enhanced vCA1– PL5 coactivation during DOWN states (Figure 4J). These observations suggest that delta spikes support memory consolidation in vCA1– PL network, as suggested in dHPC – neocortex network (Todorova and Zugaro, 2019).

Moreover, we observed that shock modulation of firing depended upon coupled regions in PL5, but not in BLA and vCA1 (Figure S8A). There were no significant differences in cue-modulation of firing across coupled cells in any behavioral sessions (Figure S8B) or systematic differences in freeze modulation across behavioral sessions in PL5 and vCA1 (Figure S8C). These results suggest that the information representation of each cell group is largely stable across sessions but differs across cell groups in a coupled region-dependent manner.

## Discussion

Recent studies have suggested that memory encoding cell ensembles in local circuits are configured before memory acquisition (Dragoi and Tonegawa, 2011; Mizuseki and Miyawaki, 2017), although post-acquisition stabilization may occur (Farooq et al., 2019; Grosmark and Buzsaki, 2016). Consistently, our results demonstrated that coactivation participating ensembles in BLA and PL5 were activated more than chance level prior to conditioning (Figure 8A). Moreover, we detected virtually no ensemble coactivations during pre-cond. NREM (Figures 2, 3 and S2). These findings suggest that elements of a given memory are instantly encoded in preconfigured cell ensembles in various brain regions, and *de-novo* inter-regional ensemble coactivations bind these elements together to form a new memory (Figure 8E). PL5 cells coupled with vCA1 showed increases in firing rates, modulation by SWRs, and delta spike participation between pre- and post-cond (Figures 8B–8D). Further, the conditioning induced activation of vCA1 ensembles coupled with PL5 (Figure 8A), suggesting that ensembles in vHPC may work as indexes of memory traces (Teyler and Rudy, 2007) embedded in prefrontal ensembles whose contributing cells became more active due to input from vHPC only after memory acquisition. Our results also imply that BLA–vCA1–PL5 ensemble triplets are pre-determined (Figure 6I) and activated in an experience dependent manner (Figures 5E, and 6C). Such multi-regional simultaneous activation may be involved in development of the network for information binding.

Accumulating evidence suggests that inter-regional interactions are coordinated by oscillatory events. Theta oscillations are prominent when animals are on the alert (Buzsáki, 2006; Mizuseki and Miyawaki, 2017), and theta–gamma coupling has a role in inter-regional communication (Adhikari et al., 2010; Colgin et al., 2009; Schomburg et al., 2014). Faster oscillations were essential for interregional ensemble coactivation during NREM (Figures 4A–4G, S4A–S4D) during which firing synchrony is higher than that during theta oscillations (Mizuseki and Buzsaki, 2014; Mizuseki and Miyawaki, 2017). SWR-associated synchronous activity should have a large impact on postsynaptic neurons; therefore, it is suitable for efficient off-line memory processing involving inter-regional communication when the brain is disengaged from environmental stimuli (Buzsaki, 2015). Further, ensemble coactivation during such a synchronous epoch may result in temporal compression of neuronal sequences, which can facilitate plastic changes in synaptic connections (Buzsaki, 2015). Indeed, recent works suggest that SWRs can trigger changes in the network activity and synaptic connections within dHPC (Miyawaki and Diba, 2016; Norimoto et al., 2018). Therefore, inter-regional recurring coactivation hosted by fast oscillations during NREM may induce further changes in the global network.

We observed only weak modulation of vCA1 firing activates by cortical DOWN states (Figure S4G), which is in line with previous reports showing that firing activates in dHPC and medial entorhinal cortex persisted during subsets of cortical DOWN states (Hahn et al., 2012; Isomura et al., 2006). Moreover, we found that vCA1–PL5 coactivations occurred during DOWN states (Figure 4J), and vCA1 coupled excitatory cells in PL5 emit delta spikes more frequently after memory acquisition (Figure 8C), similar to previous observation in dHPC–neocortex network (Todorova and Zugaro, 2019). In contrast, BLA firing activities were strongly suppressed during cortical DOWN states (Figure S4G), BLA–PL5 ensemble coactivation preferentially preceded UP–DOWN state transitions (Figure 4J), and memory acquisition did not change delta spike participation of PL5 excitatory cells that were coupled with BLA (Figure 8C). These results suggest that delta spikes selectively involved in memory consolidation mediated by cortico-hippocampal networks. In addition, vCA1–PL5 ensemble coactivations developed through memory consolidation, whereas BLA–PL5 ensemble coactivation existed at the time of memory acquisition (Figures 6 and S6). It further implies that developing *de novo* inter-regional ensemble networks, but not maintaining existing ones, might involve delta spikes, which occur when most neocortical neurons are silent, thereby preventing interference with pre-existing networks.

It has been reported that inter-regional circuits involved in fear memory shift with time, and these circuit shifts take a day to several weeks (Do-Monte et al., 2015; Kitamura et al., 2017). In contrast, BLA–PL5 and vCA1–PL5 coactivations developed rapidly (Figures 6D, 6E, and S5B), and their time courses were comparable to the time window of synaptic consolidation (range: seconds or minutes to hours (Klinzing et al., 2019)). This finding implies that BLA–PL5 and vCA1–PL5 ensemble coactivations are consequences of plastic changes in the inter-regional synaptic connections (Degenetais et al., 2003; Pérez-Jaranay and Vives, 1991), which is in line with our observation that the time-lag of ensemble coactivation congruent with mono-synaptic inter-regional projections (Figure 3D). These fast changes may, in turn, drive slower changes that support system consolidation.

It remains unclear how vCA1–PL5 coactivation can be developed without activation during memory acquisition, but BLA– vCA1–PL5 triple-activation might play a role. We observed that activation of coupled triplets were transiently activated at shock onsets (Figure 6C) and event rates of triple-activation was enhanced during conditioning sessions compared to baseline or retention sessions (Figure 6I). Although higher triple-coactivation rate of coupled triplets than that of non-coupled triplet during baseline sessions (Figure 6I) implies that ensemble triplets are pre-determined, coupled triplet activities significantly elevated during post-cond. NREM than during pre-cond. NREM (Figure 5E). Thus, it is possible that experience-dependent activation of pre-determined network of ensemble triplets may provoke further changes in inter-regional network including vCA1 and PL5. Consistent with this, BLA activation immediately after learning may facilitate memory consolidation by modulating neuronal activity outside of the amygdala (McGaugh, 2004; Paz and Pare, 2013); triple-activation can play a role in the proposed process in memory consolidation. Our finer time resolution analyses revealed that vCA1–PL5 coactivation developed after the emergence of BLA– vCA1– PL5 ensemble triple-activation and BLA–PL5 ensemble coactivation (Figures 6G–6I, S6A). Further investigation is required to clarify whether BLA– vCA1– PL5 ensemble triple-activation and/or BLA–PL5 ensemble coactivation are needed for the emergence of vCA1–PL5 ensemble coactivation or activation of BLA alone is sufficient to develop vCA1–PL5 ensemble coactivation.

The inter-regional ensemble coactivation reappeared during memory recall (Figures 6G and 6H), and the coactivations were hosted by fast oscillations in various brain regions (Figure 7). This suggests that inter-regional ensemble coactivation during fast oscillations in wakefulness support memory retrieval, which is in line with the notion that awake SWRs in the dHPC support spatial memory (Jadhav et al., 2012). In the dHPC, replays of firing sequences associated with awake SWRs occur in both forward and reverse orders (Diba and Buzsaki, 2007), and reverse replays are selectively involved in memory updates (Ambrose et al., 2016). Although it is unclear whether diversity of reactivation, such as forward and reverse replay, also exists in vHPC SWRs, HFOs, and cRipples, it is possible that only a subset of ensemble coactivations that we observed is involved in memory updates, such as extinction learning. Thus, precisely structured sequence of inter-regional coactivations, which could not be examined in this study, should be scrutinized in the future to further understand the temporal aspect of memory process.

Overall, our study suggests that *de novo* inter-regional coordination of pre-configured local ensembles form a new memory. Although our findings imply close association between memory functions and inter-regional ensemble coactivations, necessity and sufficiency of the coactivations with respect to memory functions should be elucidated by loss/gain of function studies in the future. Thus, further studies are warranted to elucidate how changes in inter-regional ensemble coactivation are involved in memory processes and how such changes are regulated.

## Acknowledgments

We thank Joshua Johansen for advice on eyelid electrical stimulation, and Sakura Okada and Nobuyoshi Matsumoto for advice on ECG recording. We are grateful to Kamran Diba, Antonio Fernandez-Ruiz, Nathaniel Kinsky, Takuma Kitanishi, Hideyuki Matsumoto, Sebastien Royer, Yuichi Takeuchi, and Brendon O. Watson for discussion and comments on the manuscript. We thank Osaka City University Research Support Platform for letting us use the microscopies. This work was supported by JSPS KAKENHI (20K06860, 20H05477, and 19H04986 for H.M. and 20H03356, 19H05225, 16H04656, and 16H01279 for K.M.), The Uehara Memorial Foundation (H.M. and K.M.), GSK Japan Research Grant (H.M.), the Takeda Science Foundation (H.M. and K.M.), Toray Science Foundation (K.M.), the Naito Foundation (H.M. and K.M.), and the Osaka City University ‘Think globally, act locally’ Research Grant for young researchers through the hometown donation fund of Osaka City (H.M.).

## Author contributions

H.M. and K.M. designed the study, H.M. carried out all experiments and analyses, and H.M. and K.M. wrote the manuscript.

## Lead Contact

Further information and requests for resources should be directed to and will be fulfilled by the Lead Contact, Kenji Mizuseki.

## Materials Availability

The materials generated in this study are available from the Lead Contact upon reasonable request.

## Data and Code Availability

The datasets supporting this study are available from the Lead Contact upon reasonable request. All codes for the manuscript are available at https://github.com/HiroMiyawaki/miyawaki_2020.

## Methods

### Animals

Fifteen male Long–Evans rats (9.6–15.0 weeks old, 330–503 g at the time of surgery; Japan SLC) were maintained in 12-h light/12-h dark cycle (light on at 8:00 a.m.). To exclude any potential effects of estrous cycles on neural activities and animal behaviors, only male rats were used. All procedures of animal care and use were approved by the Institutional Animal Care and Use Committee of Osaka City University (approved protocol #15030) and were performed in accordance with the National Institutes of Health *Guide for the Care and Use of Laboratory Animals*.

### Surgery

The rats were anesthetized with isoflurane (1–3% in 50% air/50% oxygen mixture gas); then, small incisions were made on pectus skins, and Teflon insulated stainless wires (AS636, Cooner wire) were sutured on the left intercostal muscles to record the electrocardiograms (ECGs) (Okada et al., 2016). The other ends of the wires were subcutaneously led to small incisions made on nuchal skins. After suturing the pectus skins, the rats were placed on stereotaxic frames (Model 962, Kopf), and small pieces of the scalps were removed. Two stainless wires were inserted into the nuchal muscles for each rat for electromyogram (EMG) monitoring (Miyawaki and Diba, 2016). Short conductive wires (36 AWG, Phoenix Wire) were soldered on stainless screws (B002SG89KW, Antrin), and the screws were put on the right olfactory bulb (OB; ML +0.5 mm, DV +9.0 mm from bregma) to record electroolfactogram (EOG), which reflect respiration (Chaput, 2000). Two additional screws with 36 AWG wires were put on the cerebellum through small holes made on the skulls for ground and reference of all electrophysiological recordings. Wires for ECG, EMG, and EOG were gathered on single connectors for 16-channel differential input pre-amplifiers (C3323, Intan). Two tungsten wires (100 μm in diameter, California Fine Wire) were implanted into each eyelid, and the free ends were placed on single connectors for stimulation. Three 1 mm × 2 mm rectangular craniotomies centered at (ML +1.0 to +1.5 mm, AP +2.90 to +3.25 mm), (ML +4.60 to +4.80 mm, AP −2.60 to −3.00 mm), and (ML +2.80 to +3.00 mm, AP −4.95 to −5.55 mm) were made for recording from the prefrontal cortex, amygdala, and vHPC, respectively. Silicon probes (Buzsaki64sp and Buzsaki64spL from Neuronexus or F6-64 from Cambridge Neurotech) were attached on three-dimensional printed microdrives (STL data are available at https://github.com/Mizuseki-Lab/microdrive) and then coated with poly(3,4-ethylenedioxythiophene) conducting polymer (Yang et al., 2005) by applying direct current (0.1 µA for 3 s for each channel) controlled by nanoZ impedance tester (White Matter). The probes were inserted into the brains through the craniotomies with angles of −14, 0, 14 degrees to the D-V axes for recording from the prefrontal cortex, amygdala, and vHPC, respectively. Small Faraday cages were made with copper mesh on the skulls and secured with dental cement (Orthofast, GC) to reduce electrical noise and protect the implants.

### Electrophysiological recordings

All implanted probes and the connectors hosting ECG, EMG, and EOG signals were connected to a recording system (C3100 256 ch acquisition board from Intan or 512 ch acquisition board from Open Ephys) via pre-amplifiers (C3323 or C3325, Intan). Accelerations of the head were obtained with accelerometers on the pre-amplifiers. All signals were recorded with Open Ephys GUI software available at https://open-ephys.org. Positive polarity is up throughout this paper.

### Fear conditioning

The behaviors of the rats were recorded at 25 frames/s using a video camera (CM3-U3-31S4C-CS, Flir) with 8-mm lens (LENS-80T4C, Tamron) mounted on the ceiling. Shutter timing was controlled by a stimulator (SEN-7203, Nihon Kohden), and transistor-transistor logic pulses sent from the camera were captured with the electrophysiological recording system to obtain acquisition timing of individual frames. Behavioral experiments consisted of five sessions: baseline, conditioning, context-retention test, cue-retention test/extinction, and test for retention of extinction (Figures 1C and 1D). Conditioning and context retention tests were performed in a tube (30 cm diameter, 51 cm depth) with horizontal stripes put on metal grids scented with 1% acetate. Other behavioral sessions were conducted in a rectangular box (27 cm × 33 cm, 40 cm depth) with vertical stripes put on white plastic floor scented with 70% ethanol. Thirty seconds, 5 kHz pips (250 ms on, 750 ms off, 74 dB) were used as conditioned stimuli (CS), and trains of 2 ms electrical pulse (lasting 2 s; 4.6–5.1 mA at 8 Hz for each eyelid; left and right eyelids were stimulated alternatively with half-cycle temporal shift; generated with isolators SS-202J, Nihon Kohden) applied through eyelid wires (Johansen et al., 2010) were used as unconditioned stimuli (US). There were 700–750 ms traces between offsets of the last pips and the onsets of the first shocks. Each session started with 4-min free exploration periods in which no tone was presented, and 4 CS for baseline, 12 for conditioning, 0 for the context-retention test, 40 for cue-retention test/extinction, and 8 for test for retention of extinction were then presented. CS were presented with pseudo-random interval uniformly distributed in the range of 180–240 s, except for the last 32 tones in the cue-retention test/extinction sessions where intervals were uniformly distributed in the range of 60–120 s. US were presented only in the conditioning sessions. The duration of context retention sessions was 4 mins, and other sessions ended 4 mins after offset of last CS presentation. Context retention sessions were immediately followed by cue-retention test/extinction sessions, and other sessions were separated with 2.5–2.6-hour rest/sleep sessions in the homecages (Figure 1D). Recordings of animal behavior and electrophysiological activity in rest/sleep sessions were performed for > 2.5 hours prior to baseline sessions and continued for > 2.5 hours after the test for retention of extinction. Baseline sessions started at 8:40 a.m., and test for retention of extinction sessions ended at 7:45 p.m.; all behavioral sessions and interleaved homecage sessions were during the light cycle, whereas the first and last homecage sessions were largely during the dark cycle. Right after recordings, animals were anesthetized with isoflurane (1–3% in 50% air/50% oxygen mixture gas), and electrode positions were marked by micro-lesioning by applying DC currents (3 µA for 10 s, A365, World Precision Instrument) through the electrodes on the top and bottom of each shank.

### Histological reconstruction of electrode positions

Twelve to thirty-six hours after micro-lesioning as described above, the rats were perfused with 4% paraformaldehyde (441244, Sigma-Aldrich), and the brains were removed from the skulls. After 24 to 48 hours post-fixation in 4% paraformaldehyde, the brains were sliced into 50 or 75 μm thickness with a vibratome (VT1200S, Leica). The slices were permeabilized with 0.3% Triton X-100 (35501, Nacalai Tesque) in phosphate-buffered saline (PBS) for 30 min at room temperature and stained sequentially with NeuroTrace Red fluorescent Nissl Stain Solution (200 × dilutions; N21482, Thermo Fisher) overnight at 4 °C and with DAPI (0.5 µg/ml; D1306, Thermo Fisher) for 30 min at room temperature in PBS. The slices were washed with PBS for 30 min at room temperature before and after each staining. Micrographs of the slices were obtained with a confocal microscope (LSM 700, Zeiss) or fluorescent microscope (BZ-X800, Keyence), and the position of electrodes was reconstructed by visual detection of micro-lesioned sites (Figure 1A). The reconstructed position of the electrode tips is summarized on diagrams found elsewhere (Paxinos and Watson, 2007) (Figure S1A).

### Spike sorting

To avoid potential contamination of shock artifacts, shapes of the shock artifacts estimated with the third-order Savitzky–Golay filter (5-ms window width) were subtracted from the recorded trace in periods from 10 ms prior to shock onsets to 1 s following shock offsets whose edges were attenuated exponentially (τ = 2 ms and 200 ms for onset and offset, respectively). The subtraction was performed before spike detection. Spike detection and automated clustering were performed with Kilosort2 (Stringer et al., 2019) (available at https://github.com/MouseLand/Kilosort2). To ensure that no spikes were contaminated from the shock artifacts, spikes detected around each shock pulse (0.1 ms prior to onsets to 5 ms following the offset of each 2 ms pulse: in total, 7.1 ms per shock) were discarded, and the clusters were then manually curated on phy software (available at https://github.com/cortex-lab/phy). Lastly, we evaluated cluster quality. Isolation distance (Harris et al., 2001) was calculated with clusters detected on the same shank. Inter-spike intervals (ISI) index (Fee et al., 1996) was calculated 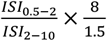, where *ISI*_x-y_ is counts of ISI in [x, y] ms window. The ISI index is useful for most cases but not for some non-bursty cells; thus, we also used the contamination rate that was introduced in Kilosort2. The contamination rate was calculated as *min* 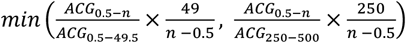, where *ACG*_x-y_ is counts of spike autocorrelogram in [x, y] ms window, and *n* is shifted from 1.5 ms to 9.5 ms with 1 ms step to find the minima. Mean waveform of each cluster was calculated on high-pass filtered (> 300 Hz) traces, a channel with maximum spike amplitude was found, and mean waveform of that channel was then upsampled to 200 KHz using spline function on MATLAB (Mathworks). Spike amplitude and spike width were determined as trough depth from the baseline and time from the trough to the peak estimated using the upsampled mean waveform, respectively. We set four criteria for cluster quality: (1) isolation distance > 15, (2) ISI index < 0.2 or contamination rate < 0.05, (3) overall mean firing rates > 0.01 Hz, and (4) spike amplitudes > 50 μV. Units that met all four criteria were used for further analyses.

### Classification of excitatory and inhibitory neurons

To classify recorded cells into excitatory and inhibitory cells, we first detected putative excitatory- and inhibitory-synaptic connections (Figure S1C) using methods described previously (Fujisawa et al., 2008) with minor modifications. For each cell pair, CCG of spike timings of two neurons was calculated in 0.1 ms bins and then smoothed with Gaussian filter (σ = 0.5 ms). The same procedure was performed on spike trains with uniformly distributed random jitters (± 5 ms range) for both cells, and maxima and minima of CCGs in the range of [−5 ms, +5 ms] were detected. The procedure was repeated 1,000 times to obtain both the mean and 99% confidence intervals of CCGs, and 99% global bands were defined as 99^th^ percentiles of maxima and minima in the range of −5 ms to +5 ms. If actual smoothed CCG had peak/trough higher/lower than upper/lower boundary of 99% global bands in [+1 ms, +4 ms] periods, the cell pair was marked as a candidate pair with monosynaptic excitatory/inhibitory connection. Smoothed CCGs of the candidate pairs were visually inspected to exclude suspicious connections such as CCG with broad peaks/troughs or strong peak/trough at time zero. The remaining pairs were accepted as pairs with monosynaptic excitation/inhibition. Cells with at least one excitatory/inhibitory innervation and no inhibitory/excitatory ones were labeled as excitatory/inhibitory cells. In all the recorded regions, excitatory cells had wider waveforms than inhibitory cells (Figure S1D). Thus, to classify cells that were not labeled as excitatory or inhibitory based on CCGs, we used spike width as previously described (Bartho et al., 2004). We labeled CCG based non-classified cells with spike width > 0.6 ms and < 0.5 ms as excitatory and inhibitory cells, respectively. CCG based non-classified cells whose spike width was between 0.5 and 0.6 ms were categorized as non-classified. The numbers of excitatory, inhibitory, and non-classified cells are summarized in Table S1.

### Sleep scoring

Sleep states were automatically scored with prefrontal LFP, hippocampal LFP, and head acceleration (Watson et al., 2016) using Buzcode scripts (available at https://github.com/buzsakilab/buzcode). The scoring results were visually inspected with the power spectrum in the prefrontal LFP, vHPC LFP, and nuchal EMG, and the scoring was modified if needed. Microarousals (Miyawaki et al., 2019; Watson et al., 2016), short (< 40 s) awake periods, which interleaved in NREM or occurred on transitions from REM to NREM, were treated as a part of NREM.

### Heart rates, EOG power spectrum, nuchal EMG amplitudes, and head accelerations

Individual heartbeats were detected as peaks on ECG signals, and mean heart rate was calculated in 0.5 s bins and smoothed using the 5-s window moving average. Multitaper power spectrum analyses of EOG signals were performed in 1-s sliding windows with 0.5-s steps using the Chronux toolbox (available at http://chronux.org). Nuchal EMG was high-pass filtered (> 10 Hz), its envelope was obtained by Hilbert transform, and amplitudes were obtained as the average of an envelope in 0.5-s bins. Accelerometer signals were high-pass filtered (> 1 Hz) on x-, y-, and z-axes signals separately to remove the effect of gravity acceleration (da Silva et al., 2018). Then, head acceleration was calculated as mean absolute values of acceleration vectors in 0.5-s bins.

### Freeze detection

Time periods of freezing behavior were detected using the Gaussian mixture hidden Markov model (HMM) with three hidden states. In the HMM, heartrate, 0.5–5 Hz, and 5–10 Hz bands power of EOG, nuchal EMG amplitudes, and the logarithm of head acceleration were used as observed variables. All behavioral sessions were concatenated; if the rat slept during behavioral sessions, such periods were excluded from detection, and HMM was then optimized for each rat. Mean head acceleration in each hidden state was calculated, and periods in the state with the slowest acceleration were labeled as freezing. Freezing periods < 5 s were removed, and freezing periods separated with gap < 1 s were concatenated.

### Wavelet analyses

Discrete wavelet transform was computed with a MATLAB wavelet software package (provided by C. Torrence and G. Compo [https://github.com/chris-torrence/wavelets]), and the wavelets power was then z-scored within each scale using means and standard deviations (SDs) within NREM sleep (Sullivan et al., 2014).

### Detection of hippocampal sharp-wave ripples

SWRs were detected on LFPs recorded in vCA1, vHPC CA3 region, or ventral subiculum using a previously described method (Miyawaki and Diba, 2016) with a minor modification. We used a slower frequency band and lower power threshold since the ripples were slower and weaker in the vHPC than in the dHPC (Patel et al., 2013). To exclude gamma contamination, we adopted ripples co-occurring with sharp-waves (Fernandez-Ruiz et al., 2019). First, for each channel, the LFPs were bandpass filtered (100–250 Hz), and their root mean squares (RMS) in 13.3 ms windows were z-scored using means and SDs within NREM sleep. Periods with signals > 1.5 z were used as candidate events on each channel. Candidates with peaks < 4 z or shorter < 30 ms were discarded, and those separated with < 10 ms intervals were concatenated. The maxima of z-scored smoothed ripple power within the candidate and its corresponding time were considered ripple peak amplitudes and the ripple peak time of the candidate, respectively. Overlapped ripple candidates detected on the same shank were concatenated, and then, candidates > 750 ms were discarded. Sharp-waves were detected using the difference between the top and bottom channels within each shank. The difference was bandpass filtered (2–40 Hz) and z-scored with mean and SD within NREM sleeps, and periods in which the signals were < −2.5 z for > 20 ms were accepted as sharp-waves. Sharp-waves > 400 ms were discarded, and sharp-wave troughs were determined as timepoints with minima of the filtered signal. If no sharp-wave troughs were detected on the same shank during ripple candidates, the candidates were discarded. Moreover, overlapped candidates detected on different shanks were concatenated, and events > 750 ms were then discarded. In the case where candidates detected on multiple channels/shanks were concatenated, the ripple peak time of the candidate with maximum ripple peak amplitude (in z-score) was used as the ripple peak time of the concatenated candidate, and the corresponding channel was regarded as the maximum ripple power channel for the SWRs. Otherwise, a channel on which the interested candidate was detected was regarded as the maximum ripple power channel of the SWRs. For each SWRs, ripple troughs on the maximum ripple power channel were detected, and the SWR peak time was defined as the time point of the ripple trough closest to its ripple peak time.

### Detection of amygdalar high-frequency oscillations

HFOs in the amygdala (Ponomarenko et al., 2003) were detected as previously described for amygdalar high-gamma detection (Amir et al., 2018) with a minor modification. The median of LFPs for each shank was bandpass filtered (90–180 Hz), and the RMS of the filtered signals (window = 20 ms) were then converted to z-score using mean and SD within NREM sleeps. Periods with z-score > 2 were classified as candidate events. Candidates with peaks < 4 z were discarded, candidates with < 30 ms duration were removed, then candidates separated with intervals < 20 ms were concatenated. Detection was performed for each shank. The peak time of an HFO candidate was determined as the time of peak on the square-roots of the smoothed bandpass filtered signals used for the detection, and its peak height (in z-score) was used as the peak power. Overlapped candidate events on different shanks were concatenated, and events > 750 ms were discarded. When the HFO candidates detected by multiple shanks were concatenated, the highest peak power (in z-score) across shanks and the corresponding peak time were defined as peak power and time of the concatenated candidate, respectively. Because HFOs occur mainly during NREM (Ponomarenko et al., 2003), we accepted only candidates detected during NREM periods as HFOs, and candidates within awake periods were classified as aHFOs. To estimate peak frequency of HFO, wavelet power was calculated on the bandpass filtered median LFP of the shank with maximum peak power determined as described above for each HFO (in 90–180 Hz band, 11 scales). The wavelet power peak was then detected, and its corresponding frequency was assigned as the peak frequency of the interested HFO.

### Detection of prefrontal gamma and ripple oscillations

Prefrontal γ_slow_, γ_fast_, and cRipple oscillations (Khodagholy et al., 2017) were detected on LFPs in the prelimbic cortex. The LFPs of each channel were bandpass filtered (30–60 Hz, 60–90 Hz, and 90– 180 Hz for γ_slow_, γ_fast,_ and cRipples, respectively), and RMS (window = 20 ms) of the filtered signal were z-scored with mean and SD within NREM epochs. Periods with z-score > 3 were used as candidate events, and the maxima of the RMS within the candidate and its corresponding time points were classified as the peak power and the peak time of the candidate, respectively. Candidates with peak power < 5 z were discarded, candidates with duration < 50 ms were removed, then candidates separated by < 30 ms intervals were concatenated. Detection was done for each channel separately; then, overlapped events on different channels were concatenated, and events > 750 ms were discarded. When concatenating multiple candidates, the peak time corresponding to the highest peak power (in z-score) across channels was assigned as the peak time of the oscillatory events.

### Detection of prefrontal sleep spindles

Spindles in the PL were detected on wavelet power, as described previously (Sullivan et al., 2014). Because spindles were recorded in multiple channels/shanks coherently, wavelet transform was performed on mean LFP across all PL cortical channels (in 9–18 Hz band, 11 scales) and was normalized as described above (see ‘Wavelet analyses’ for details), and the maximum on the normalized wavelet power across the calculated scales was obtained in each time point. Epochs with the maximum normalized wavelet power > 1.4 z for > 350 ms within NREM epochs were marked as candidate spindles. When peaks of the maximum normalized wavelet power within candidates were < 2 z, the candidates were discarded. For each spindle event, the time point with the maximum normalized wavelet power was marked as peak time.

### Detection of prefrontal delta waves and delta spikes

Mean LFP across channels in the PL were calculated, and delta waves were detected as positive deflections of bandpass (0.5–6 Hz) filtered mean LFPs during NREM (Maingret et al., 2016; Miyawaki and Diba, 2016; Vyazovskiy et al., 2009). Sequences of upward-downward zero crossings of the z-scored bandpass filtered signal were detected as onsets and offsets of candidate of delta waves, and peaks were taken as maximum deflection between the onsets and offsets. If deflections of the filtered signal at the peaks were < 1.5 z or durations from the onsets to the offsets were < 100 ms or > 1s, these candidates were discarded. We accepted candidates as delta waves only when the filtered signal was monotonically increasing from onset to peak and monotonically decreasing from peak to offset. Typical duration from onsets to peaks, peaks to offsets, and offsets to the next peaks were calculated as median of these intervals within individual rats. For calculation of typical duration from delta offsets to the next delta peak, we excluded events duration from delta offsets to the next delta onsets > 4 sec. We did not consider spiking activity to detect delta waves since the detection heavily depended upon the number of simultaneously recorded neurons. Furthermore, a small fraction of neurons fire even during delta peaks and are known as delta spikes (Todorova and Zugaro, 2019). We defined delta spikes as spikes occurring within ± 15 ms of delta peaks and the calculated delta spike participation rate as a fraction of delta waves in which a given cell had at least one delta spike.

### Ensemble detection with independent component analyses

Instantaneous activation strength of cell ensembles was estimated using ICA, as described previously (Giri et al., 2019; Lopes-dos-Santos et al., 2013; Todorova and Zugaro, 2019; Trouche et al., 2016). First, for each brain regions, the z-scored firing rate matrix of recorded neurons was obtained in 20-ms bins. Next, principal components analyses (PCA) were performed on it, and significant components with corresponding eigenvalues exceeding the Mar-cenko-Pastur threshold (Peyrache et al., 2010; Peyrache et al., 2009) were found. Then, the firing rate matrix was projected onto the significant components, and ICA was performed on the projected matrix using FastICA package for Matlab (available at https://research.ics.aalto.fi/ica/fastica/). The projection vector of each ensemble was calculated as products of the PCA projection matrix and the weight vector of ICA, and the projection vector was then normalized to unity length. On the z-scored firing rate matrix in a matched epoch **M**, instantaneous ensemble activation strength at timepoint t, *A*_*k*_(*t*), was calculated as **M**(*t*)^T^**P**_k_**M**(*t*). Here, **P**_k_ is the outer products of the normalized projection vector with its diagonal set to zero. We used activities during conditioning sessions as templates unless otherwise specified. We performed these analyses in each brain region separately. The numbers of detected ensembles are summarized in Table S2.

### Ensemble coactivation

To evaluate coactivation of cell ensembles, CCGs between instantaneous ensemble activation strengths were calculated as

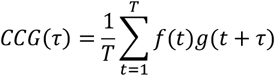

where *T* is the number of the analyzed bin, and *f(t)* and *g(t)* are z-scored instantaneous ensemble activation strengths at time *t*. Since the signals were z-scored, *CCG(τ)* gives the Pearson correlation coefficient between the signal *f* and time-shifted signal *g*. CCGs were smoothed with Gaussian kernel (σ = 20 ms) for presentation. Then, the maximum deflection of each CCG was detected within ± 100 ms window around time 0. The significance of the deflection was evaluated based on chunk shuffling described as follows. First, one of the signal pairs was divided into 2-sec chunks, the order of the chunks was randomly shuffled, and CCG was then calculated. This method preserves the finer (< 2 s) structure of auto-correlograms (ACGs), which is important because non-uniformity of ACGs may be inherited to CCG (Dean and Dunsmuir, 2016). The shuffling was iterated 500 times, and 99% confidence intervals of the maxima and minima of the CCG in a range of ± 100 ms were then estimated. The CCG peak/trough was found as a point when absolute values of differences between the actual CCG and shuffled mean reached the maximum. If the actual CCG value of the peak/trough was larger or smaller than the 99% confidence intervals, the peak or trough was regarded as significant. Pairs with significant peaks and troughs during post-cond. NREM were labeled as coupled and inverse-coupled pairs, respectively. Ensemble pairs that did not have significant peaks during post-cond. NREM were labeled as non-coupled pairs, which include inverse-coupled pairs unless stated otherwise.

To assess the contribution of SWRs, HFOs, cRipples, or spindles on the coactivations of coupled ensemble pairs, we calculated event-excluded CCGs between instantaneous ensemble activation strength of ensembles by removing time bins which contained the interested events from the CCG calculation. To investigate whether significant coactivation occurred outside of the interested oscillatory events, chunk shuffling (500 times), as described above, was performed on the event-excluded CCGs to test whether significant peaks were lost. To test whether the coactivation occurred preferentially within interested oscillatory events, we calculated peak drops as differences of peak height between event-excluded CCGs and CCGs with entire-NREM, and their significances were tested as follows. First, we performed the surrogate event-excluded CCG analysis with randomly jittered events (jitters distributed uniformly in ranges of ± 500 to ± 2,500 ms for SWRs, HFOs, and cRipples and ± 3 to ± 5 s for spindles) 500 times for each event type, and the peak drop of each CCG was calculated. Then, we tested whether the actual peak drops were larger than 99.5^th^ percentiles of peak drops in jittered events.

### Triple-activation of ensembles across brain region

To evaluate simultaneous reactivation across BLA, vCA1 and PL5 ensembles, we expanded CCG analyses for ensemble triplets by using a similar method previously used for spike triplet analyses (Nadasdy et al., 1999). First, we define triple-CCG as

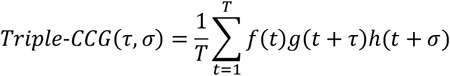

where *T* is the number of the analyzed bin, and *f(t), g(t)*, and *h(t)* are z-scored instantaneous ensemble activation strengths at time *t*. Triple-CCGs were smoothed with 2-dimensional Gaussian kernel (σ = 10 ms) for presentation. The maxima of triple-CCG and corresponding time gaps in the range of |τ|, |σ|, |τ-σ| < 100 ms, which corresponds to triple-activation occurred within 100 ms time gaps, were took as peak values and positions, respectively. To examine significance of peak values, we performed the chunk shuffling used for CCG analyses for the all three signals to generate 500 surrogate data. When the peak value of actual data was larger than top 0.5 percentile of surrogate data, the triplet was labeled as a coupled triplet.

Triple-CCG defined above can be interpreted as *h(t)* weighted CCG between *f(t)* and *g(t)* (If h(t) is uniform, the triple-CCG is identical to CCG between *f(t)* and *g(t)* up to scales). Since *h(t)* is z-scored instantaneous activation strength which stays near zero for most time (Figure 2B), most time points have minor contribution to the weighted CCG. Thus, in case that coactivation between *f(t)* and *g(t)* occurred only when *h(t)* is far above zero, getting triple-CCG improve signal-to-noise ratio of CCG between f(t) and g(t). Because of this, even when all participated ensemble pairs do not have significant CCG peaks, the triplets may have significant peak.

### Detection of activation, coactivation, and triple-activation timings of the ensembles

The instantaneous activation strength of each ensemble was z-scored; then, activation events were detected as peaks > 5 z, and the peak strength and peak time were assigned as peak height and timestamp of the ensemble activation events, respectively. To examine significance of activation event rates, we calculated instantaneous activation strengths of surrogate ensembles (500 surrogates for each ensemble) obtained by randomly permutation of ICA projection vector. Then activation events were detected on them as peaks > 5 z, and their event rates were obtained during pre- and post-cond. NREM.

To detect timepoints of coactivations, first, the optimal temporal shift of each ensemble pair was determined by detecting peak timing on CCG of the instantaneous activation strength of the pair during post-cond. NREM. Next, we calculated instantaneous coactivation strength as the product of z-scored instantaneous ensemble activation strength with the optimal temporal shift. Coactivation events were detected as peaks > 25 z^2^ on the instantaneous coactivation strength. For each coactivation event, maxima of the coactivation strength and its corresponding time were classified as coactivation peak height and timestamp of the coactivation events, respectively. Similarly, instantaneous triple-activation strength was calculated as the product of time-shifted activation strength whose shifts were determined based on the triple-CCG peak position during post-cond. NREM, then triple-activation events were detected as peaks > 125 z^3^ on it. For each triple-activation event, maxima of the triple-activation strength and its corresponding time were classified as its peak height and timestamp, respectively.

Time gaps between co-activated and triple-activated ensembles are not necessarily same during sleep and awake periods. Thus, for analyses of coactivation/triple-activation during behavioral sessions, we re-calculate optimal time shift using CCGs/triple-CCGs within interested behavioral sessions, then coactivation/triple-activation events were detected as described above.

Shock triggered histograms of the coactivation/activation occurrence rate and mean peak height were calculated in 20 ms bins and then smoothed with Gaussian kernel (σ = 20 ms). In the homecage sessions preceding and following the conditioning sessions, the occurrence rate and mean peak heights of activation/coactivation events were calculated in 3 min bins. Those of triple-activation events were calculated in wider (10 min) bins because of their low event rates. To average activation/coactivation dynamics across animals, only NREM periods within each time bin were used to exclude effects caused by the sleep state difference.

### Coactivation triggered average of wavelet power

We selected one channel for each probe, performed wavelet transform on the LFP in 0.5–330 Hz band (94 scales), and normalized wavelet transforms as described above (see ‘Wavelet analyses’). Channels with maximum theta (6–10 Hz) and delta (0.5–4 Hz) power were used for hippocampal and prefrontal wavelet analyses, respectively. For amygdalar wavelet analyses, the channel with maximum power in gamma band (50–100 Hz) among BLA channels was used. If no BLA channels were available, lateral amygdala channels were used instead. Coactivation event-triggered average of wavelet power was calculated for each coupled ensemble pair and was then averaged within each region pair.

### Oscillatory event-triggered average of coactivation/activation strength

SWR-, HFO/aHFO-, spindle-, PL delta-, PL γ_slow_- PL γ_fast_-, and PL cRipple-peak triggered average of coactivation was calculated as a peri-event triggered average of instantaneous coactivation strength. The peri-event triggered average was calculated in ± 2 s window and then z-scored for visualization. Peaks of SWR, HFO, and cRipple triggered average of coactivation strength were detected in range of [−100 ms, +100 ms] from the oscillatory event peaks. Peaks of PL delta-peak triggered average of BLA–PL5 and vCA1–PL5 ensemble coactivation strength were detected in range of [−400 ms, 0 ms] and [−200 ms, +200 ms] from PL delta peaks, respectively.

Similarly, SWR-, HFO-, and cRipple-peak triggered average of activation was calculated as the peri-event triggered average of instantaneous ensemble activation strength. Peri-event triggered average was calculated in ± 2 s window for pre- and post-cond. NREM. For visualization, the results were z-scored within each ensemble using mean and SD across pre- and post-cond. results and plotted in range of ± 400 ms.

Peak times of SWRs and HFO were used for calculation of PL delta-peak triggered average of SWR and HFO event rates, and the event rate peaks were detected in range of [−400 ms, 0 ms] from PL delta peaks.

### Partner region of ensembles and cells

If an ensemble participated in at least one coupled inter-regional ensemble pair, the ensemble was regarded as a coupled ensemble, and its coupled region was determined as a region from which the partner ensemble was identified. For each ensemble, cells with an absolute weight of projection vector > 0.3 were defined as highly contributing cells of the ensemble. Highly contributing cells of coupled ensembles were labeled as coactivation contributing cells, and the region of the partner ensemble was defined as the coupled region of the coactivation contributing cells. Note that a given cell can be highly contributing for multiple ensembles and may have more than one coupled region.

### Modulation of cell firing

Since firing rates of individual neurons are typically log-normally distributed (Buzsaki and Mizuseki, 2014; Mizuseki and Buzsaki, 2013), mean and SD of firing rates were calculated on logarithm of firing rates to compare firing rates across cell populations and behavioral states. For analyses taking ratio of firing rates such as firing rate modulation and modulation index, mean firing rates were calculated in linear scale to avoid negative values.

The firing rate modulation by SWRs and HFOs was measured as 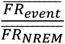,where 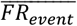 and 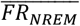 are mean firing rates within interested events (SWRS or HFOS) and entire NREM, respectively.

Modulation indices of firing rates were calculated as 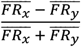 where 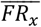 and 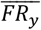 are mean firing rates in periods x and y, respectively. For the freeze modulation index, freezing and non-freezing awake periods were used as period x and y, respectively. For the shock modulation index, 0–2 s prior to shock onsets and 0–2 s following shock onsets were used as period x and y, respectively. For the cue modulation index, 0–2 s prior to cue onsets and 0–2 s following cue onsets were used as period x and y, respectively.

## Supplemental Information

**Figure S1.**
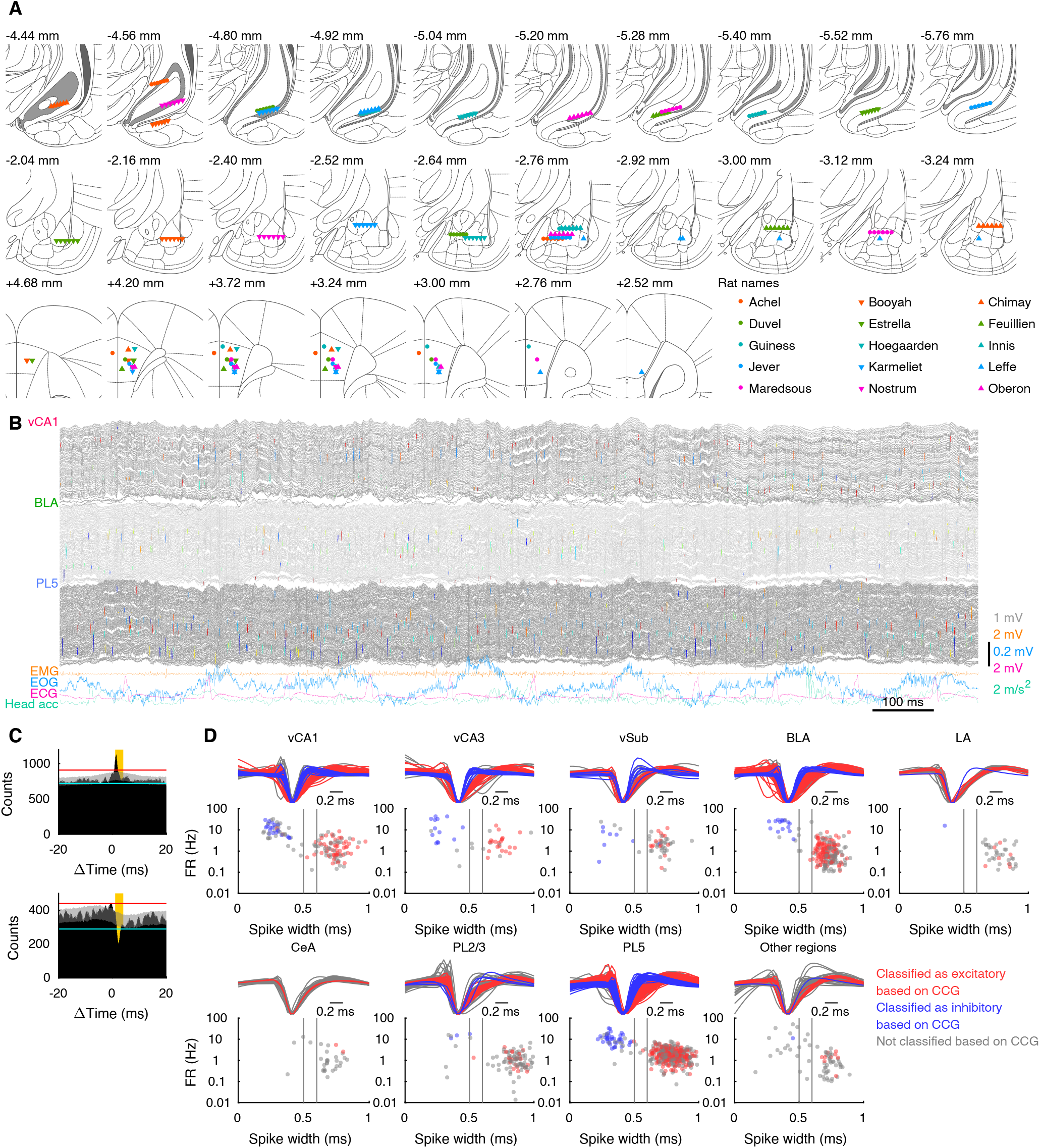
Details of electrophysiological recordings, related to Figure 1. **(A)** Positions of probe tips of all rats marked on the coronal sections of the brain atlas adapted from a previous publication (Paxinos and Watson, 2007) with permission. AP-axis coordinate from the bregma is shown at the top-left of each image. **(B)** Example traces obtained from a rat shown in Figures 1A and 1B. Traces recorded from vCA1/BLA/PL5 are shown in grey, and well-isolated units are highlighted with color. Different units are represented in different colors within each region. The bottom four colored traces are EMG, EOG, ECG, and head acceleration (Head acc). **(C)** Example CCGs of spike times with significant spike transmission (top) and suppression (bottom). Grey bands illustrate 99% confidence intervals of jittered CCG. Horizontal lines indicate upper and lower global bands. Orange background shows the period of 1–4 ms, in which the significance of peaks or troughs were evaluated. **(D)** Mean waveforms normalized with the spike amplitudes (top) and scatter plots of spike width versus mean firing rates (bottom) of all recorded units. Colors of trace/dot indicate unit classification based on CCG. Vertical lines on the scatter plots indicate thresholds for excitatory (0.6 ms) and inhibitory (0.5 ms) cells, respectively. The numbers of cells are summarized in Table S1.

**Figure S2.**
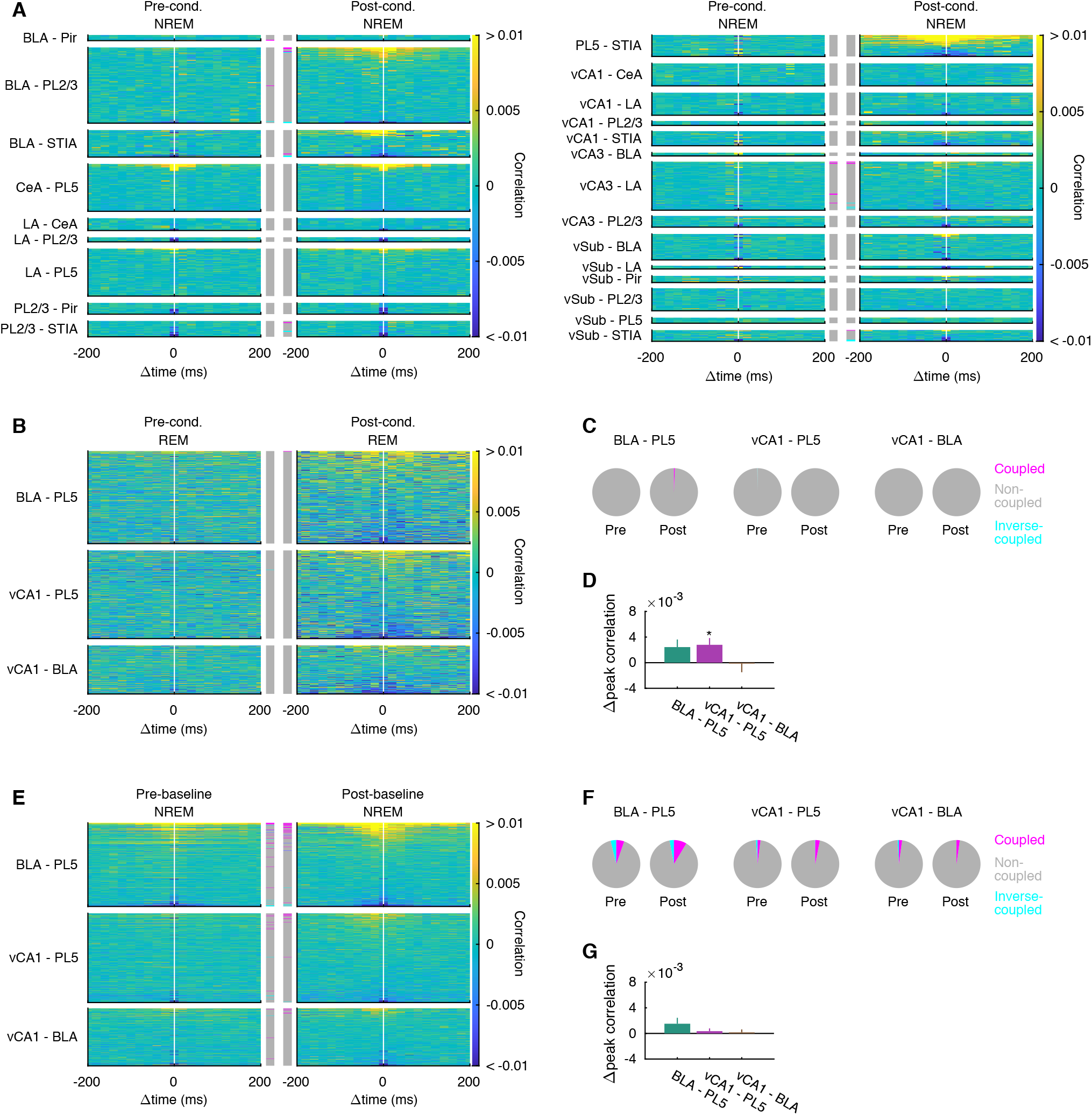
Coactivation of ensembles in various brain regions, related to Figure 3. **(A)** Inter-regional CCGs of instantaneous activation strength of ensembles identified in conditioning sessions during pre- and post-cond. NREM. Colored bars in the middle columns indicate ensemble pairs with significant peak (magenta) and trough (cyan), respectively (p < 0.01, chunk shuffling). The panels illustrate all analyzed pairs other than pairs among vCA1/BLA/PL5 shown in Figure 3A. CeA, central amygdaloid nucleus; LA, lateral amygdala; Pir, pyriform cortex; PL2/3, prelimbic cortex layer 2/3; STIA, bed nucleus of the stria terminalis intra-amygdaloid division; vCA3, ventral hippocampus CA3 region; vSub, ventral subiculum. **(B)** CCGs of instantaneous activation strength of ensembles identified in conditioning sessions during pre- and post-cond. REM sleep. **(C)** The fraction of ensemble pairs with significant peaks or troughs of CCGs shown in (B). No significant changes were detected (p > 0.05, χ^2^ test). **(D)** Changes in peak height of CCGs shown in (B). Error bars indicate SE. * p < 0.05, WSR-test. **(E-G)** CCGs of instantaneous activation strength of ensembles identified in baseline sessions during NREM in preceding and following homecage sessions (E), fraction of ensemble pairs with significant peaks or troughs (F), and changes in peak height (G) of CCGs shown in (E) (n = 338, 356, 229 pairs for BLA–PL5, vCA1–PL5, and vCA1–BLA, respectively). No significant changes are detected on (F) and (G) (p > 0.05, χ^2^ test or WSR-test). The numbers of analyzed pairs in (A–D) are summarized in Table S3.

**Figure S3.**
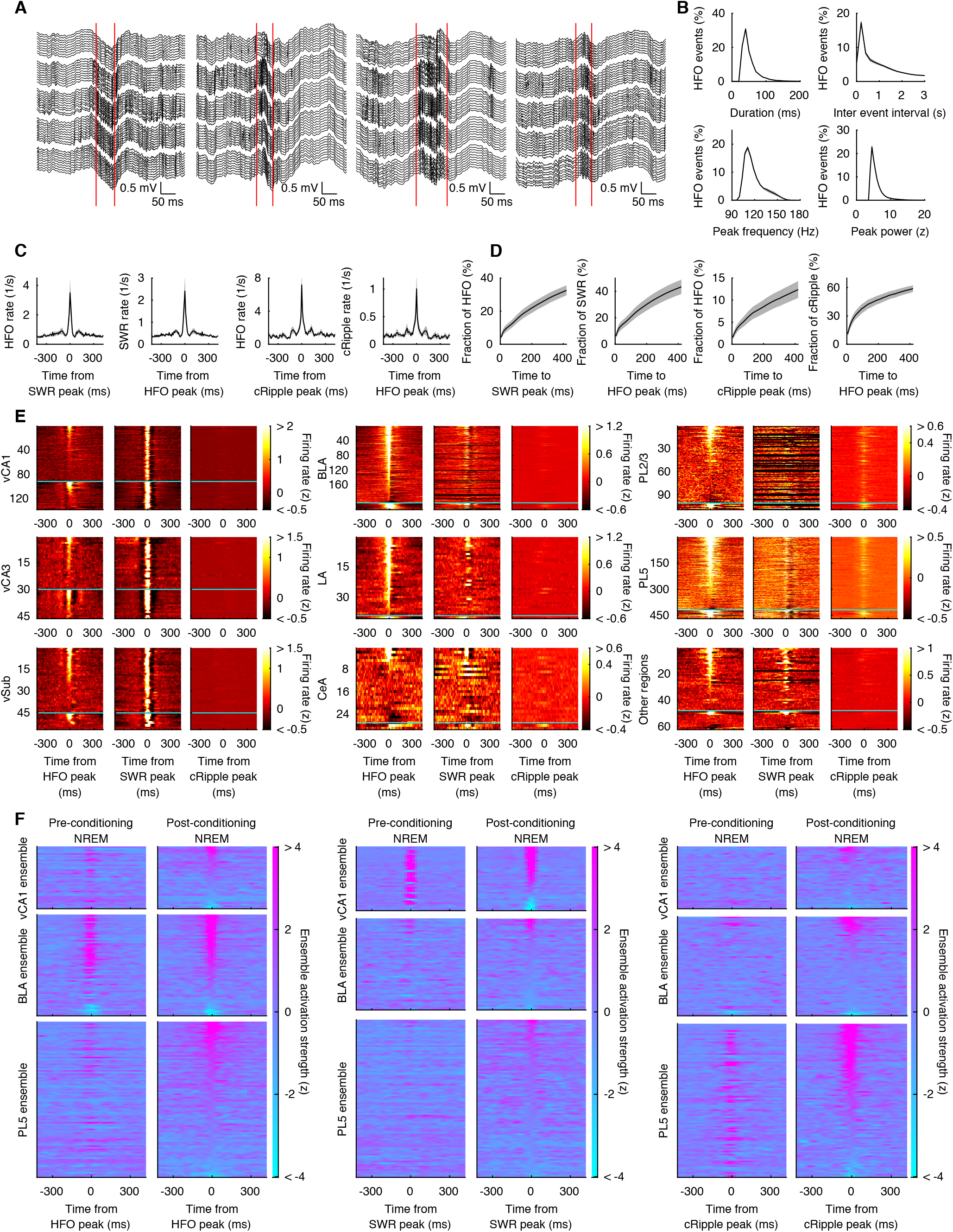
Amygdalar high-frequency oscillations, related to Figure 4. **(A)** Representative examples of amygdalar HFOs. The onset and offset of each event are marked as red vertical lines. **(B)** Histograms of duration, inter-event interval, peak frequency, and peak power of HFO. Grey sheds indicate SE (n = 15 rats). **(C)** CCGs of HFOs versus other oscillatory events. CCGs between HFO- and SWR/cRipple-peaks have a noticeable peak at time zero. Grey sheds indicate SE (n = 15 rats). **(D)** Cumulative histograms of intervals from HFO peaks to the closest SWR peaks (left), from SWR peaks to the closest HFO peaks (second left), from HFO peaks to the closest cRipple peaks (second right), and from cRipple peaks to the closest HFO peaks (right). Grey sheds indicate SE (n = 15 rats). **(E)** HFO-, SWR-, and cRipple-peak triggered histograms of firing in each brain region. Each row indicates each cell. Cells plotted above horizontal cyan lines are excitatory cells, and the remaining are inhibitory cells. Cells were ordered based on peak height at time zero of HFO triggered histograms. The numbers of cells are summarized in Table S1. **(F)** HFO-, SWR-, and cRipple-peak triggered histograms of instantaneous ensemble activation strength in pre- and post-cond. NREM. Ensembles were ordered based on peak height at time zero in post-cond. sessions for each trigger type. The numbers of ensembles are summarized in Table S2.

**Figure S4.**
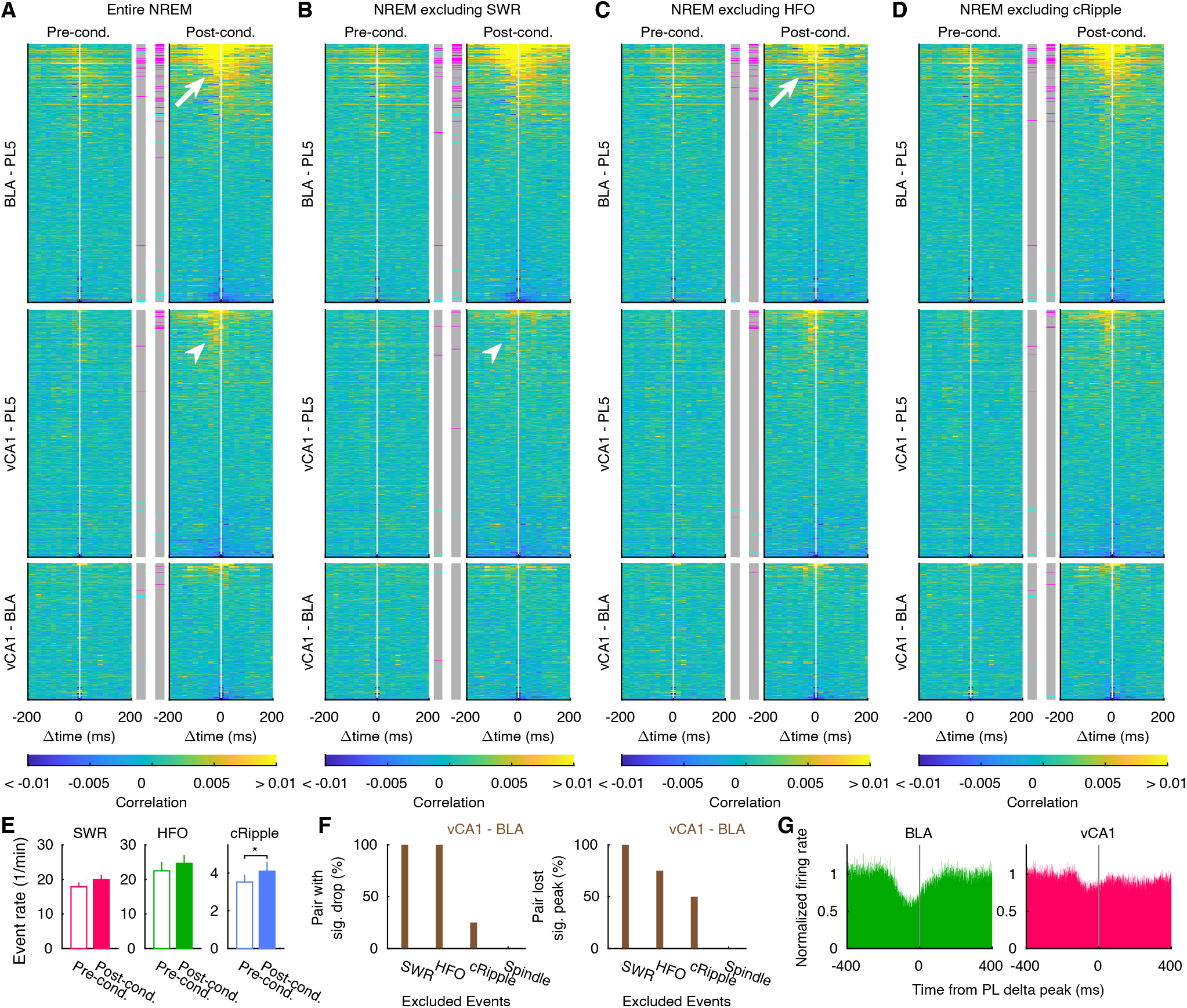
Inter-regional coactivation outside of hippocampal SWRs and amygdalar HFOs, related to Figure 4. **(A–D)** CCGs of instantaneous ensemble activation strength in entire NREM (A), NREM outside of SWRs (B), NREM outside of HFOs (C), and NREM outside of cRipples (D). CCGs in entire NREM (A) are identical to those in Figure 3A and are presented here just for comparison. Ensemble pairs are sorted based on the peak height of post-cond. entire NREM CCG. White arrows/arrowheads indicate CCG peaks between BLA–PL5 / vCA1–PL5 ensemble pairs, which are less prominent on HFO-/SWR-excluded CCGs, respectively. **(E)** Mean and SE of SWR, HFO, and cRipple occurrence rates during pre- and post-cond. NREM. * p < 0.05, WSR-test (n = 14, 15, and 15 rats for SWRs, HFOs, and cRipples, respectively) **(F)** Fractions of vCA1–BLA coupled ensemble pairs that significantly reduced CCG peak height by excluding bins containing SWRs, HFOs, cRipples, or spindles (left) and those that lost significant peaks when bins containing SWRs, HFOs, cRipples, or spindles were excluded (right). Significance was evaluated based on random jittering (left; p < 0.01) and chunk shuffling (right; p < 0.01), respectively. **(G)** PL delta peak triggered average of normalized firing rates in BLA (n = 12 rats) and vCA1 (n= 8 rats). The numbers of analyzed pairs in A–E are summarized in Table S3.

**Figure S5.**
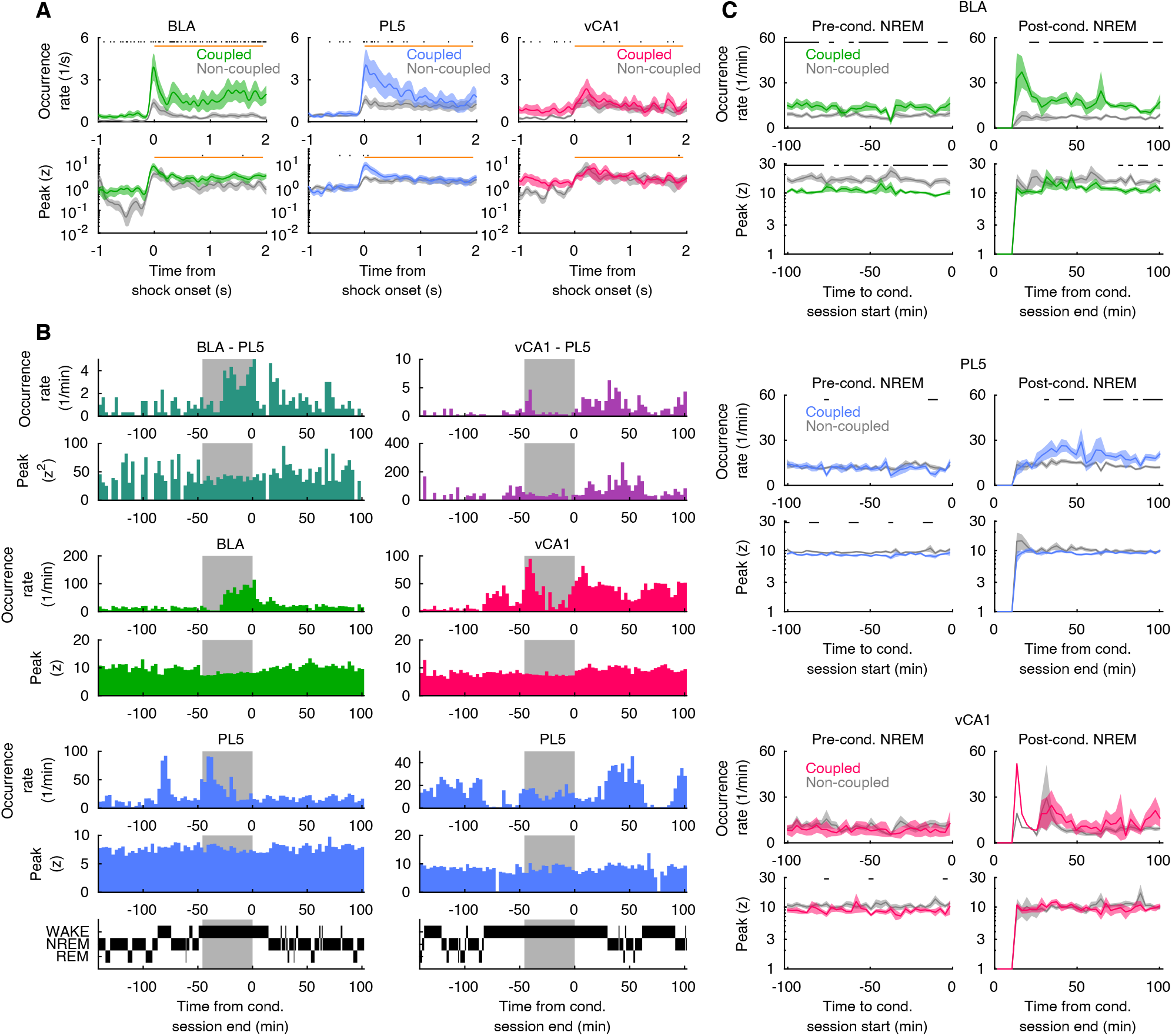
Time evolution of ensemble activation, related to Figure 6. **(A)** Shock triggered average of occurrence rates and peak height of ensemble activation events (n = 16/44, 25/68, and 10/28 coupled/non-coupled ensembles in BLA, PL5, and vCA1 respectively). Lines and Sheds represent means and SEs. Periods with a significant difference between coupled and non-coupled ensembles are indicated with black ticks on the top (p < 0.05, WSR-test). **(B)** Representative examples of time evolution in BLA–PL5 and vCA1–PL5 ensemble coactivation (top 2 rows) and activations of ensembles consisting of the coactivations (middle 4 rows). Occurrence rate and mean peak strength were plotted regardless of the brain state. Grey backgrounds indicate periods of conditioning sessions. Hypnograms are shown on the bottom. **(C)** Mean occurrence rates and peak height of ensemble activations during NREM aligned to conditioning session onset or offset (n = 16/44, 25/68, and 10/28 coupled/non-coupled ensembles in BLA, PL5, and vCA1, respectively). Lines and sheds represent means and SEs. Periods with significant differences between coupled and non-coupled ensembles are indicated with black ticks on the top (p < 0.05, WSR-test).

**Figure S6.**
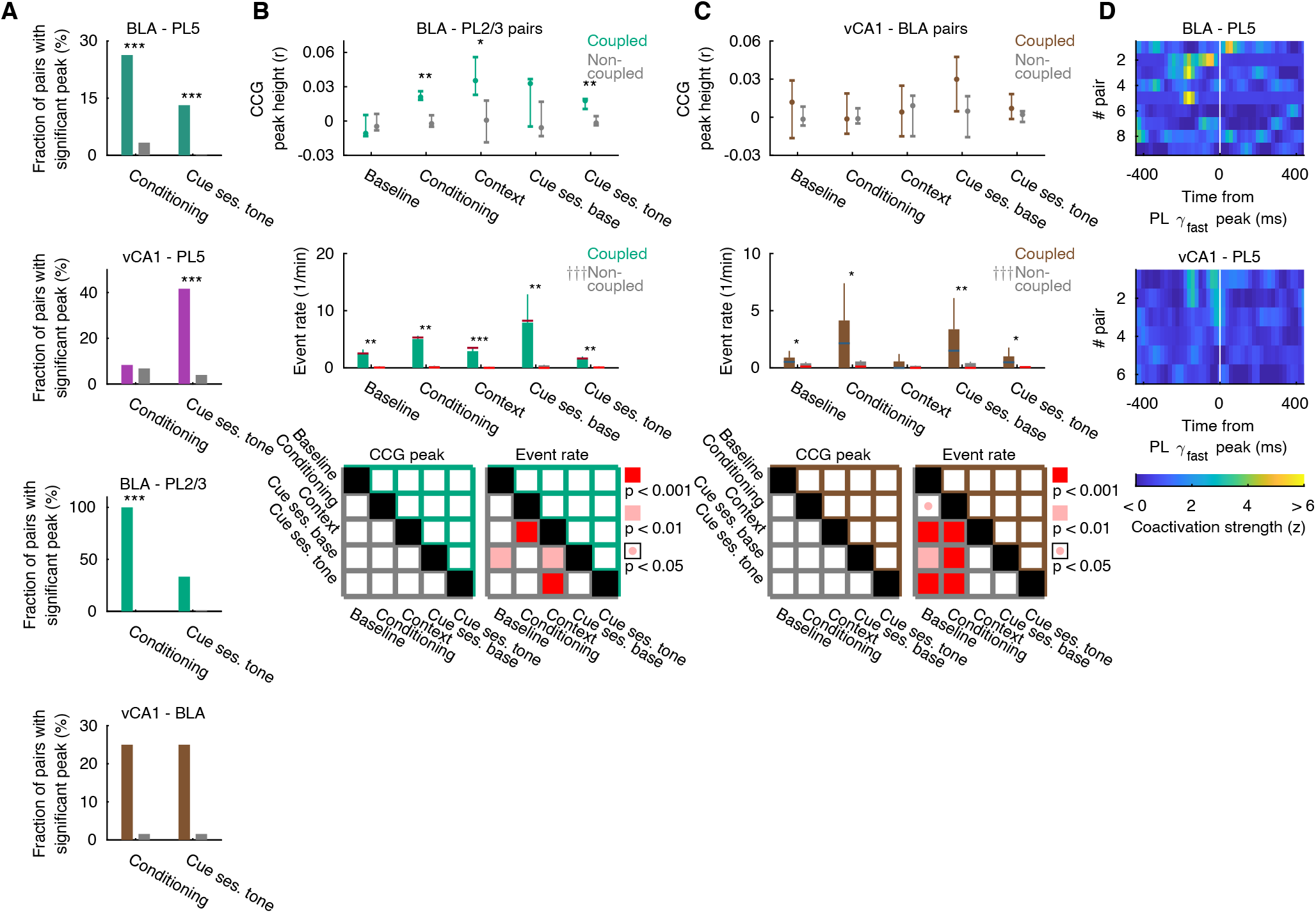
Inter-regional ensemble coactivation during behavioral sessions, related to Figures 6 and 7. **(A)** Fraction of BLA–PL5, vCA1–PL5, BLA–PL layer 2/3 (PL2/3), and vCA1–BLA ensemble pairs that showed significant peaks during conditioning and cue sessions after the first tone. Analyzed duration was matched for both sessions (45.8 min). *** p < 0.001 (Fisher’s exact test). **(B, C)** Median of CCG peak height and coactivation event rates in BLA–PL2/3 (B) and vCA1–BLA (C) ensemble pairs. Cue retention sessions (cue ses.) were divided into two parts at the onsets of the first tone. Error bars indicate IQR on the top panels and SE on the middle panels. Horizontal bars on the middle panels show medians. *** p < 0.001, ** p < 0.01, * p < 0.05 (WRS-test). ††† p < 0.001, †† p <0.01, † p < 0.05, Friedman test. Significance of pair-wise comparison across behavioral sessions are summarized on bottom (post-hoc WSR-test with Bonferroni correction following Friedman test). Top and bottom halves represent coupled and non-coupled pairs/triplets, respectively. **(D)** PL γfast-peak triggered an average of instantaneous coactivation strength during cue retention sessions. Only coupled ensemble pairs with a significant peak during cue retention sessions are shown (n = 9 / 6 ensemble pairs for BLA–PL5 / vCA1–PL5, respectively). The numbers of analyzed pairs in (A-C) are presented in Table S3.

**Figure S7.**
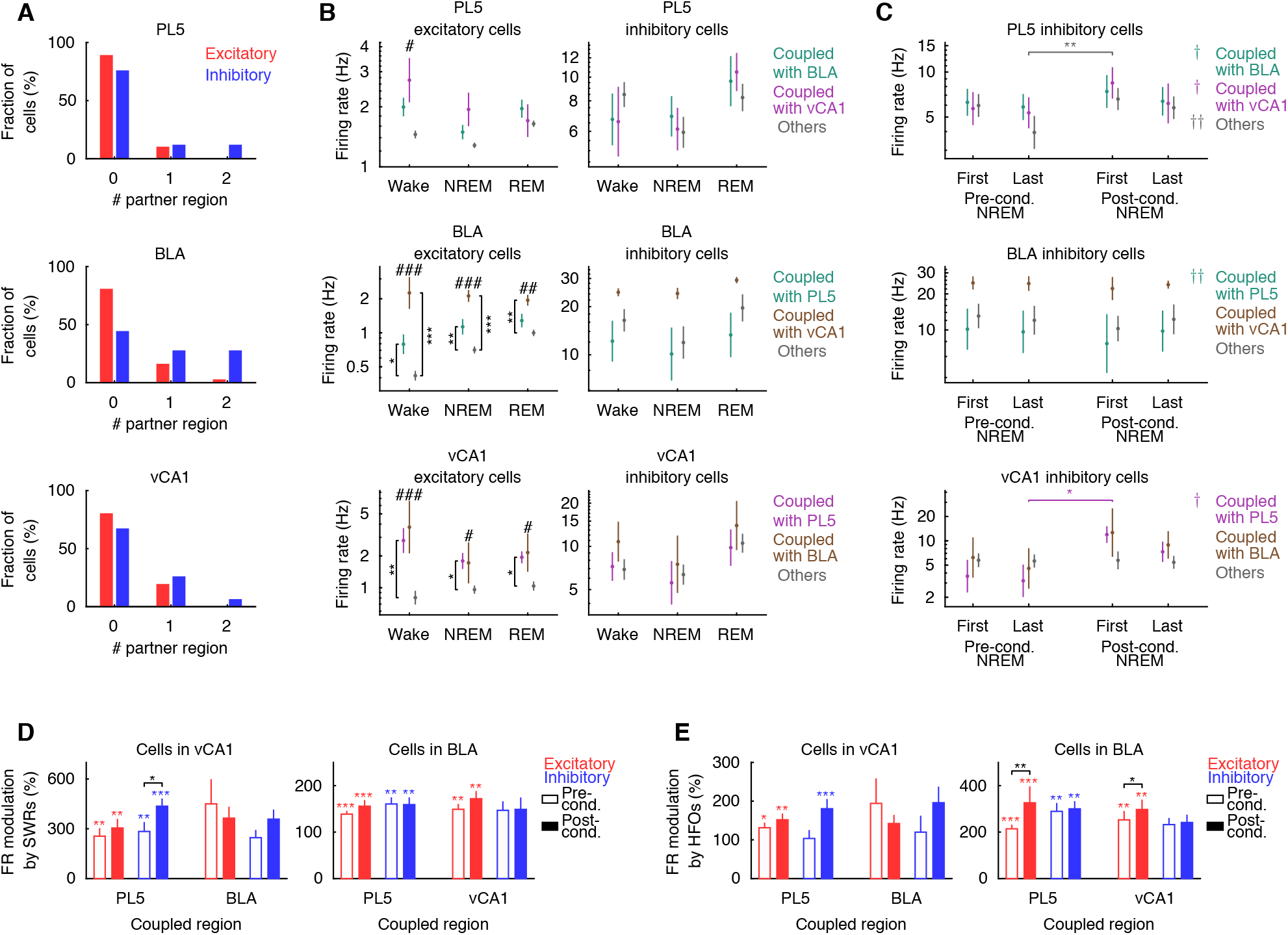
Firing properties of cells with inter-regional partners, related to Figure 8. **(A)** Histograms showing the number of paired regions per cell in PL5, BLA, and vCA1. **(B)** Mean firing rates of coactivation contributing excitatory and inhibitory cells coupled with other brain regions during wake, NREM, and REM sleep in homecage sessions. Cells that were not coupled with PL5, BLA, or vCA1 are shown as others. # p < 0.05, ## p < 0.01, ### p < 0.001, Kruskal–Wallis test; p < 0.05, ** p < 0.01, *** p < 0.001, post-hoc WRS-test with Bonferroni correction. Error bars indicate SE. **(C)** Mean firing rates of inhibitory cells during NREM in earlier and later halves of pre- and post-cond. homecage sessions. † p <0.05, †† p < 0.01, ††† p < 0.001, Friedman test; * p < 0.05, ** p < 0.01, *** p < 0.001, post-hoc WSR-test with Bonferroni correction. Error bars indicate SE. **(D, E)** Firing rate (FR) modulations by SWRs (D) and HFOs (E) during pre- and post-cond. NREM in vCA1 and BLA. Error bars indicate SE. * p < 0.05, ** p < 0.01, *** p < 0.01 WSR-test for modulation within pre- or post-cond. NREM (colored) and changes from pre- to post-cond. NREM (black). The numbers of analyzed cells are summarized in Table S6.

**Figure S8.**
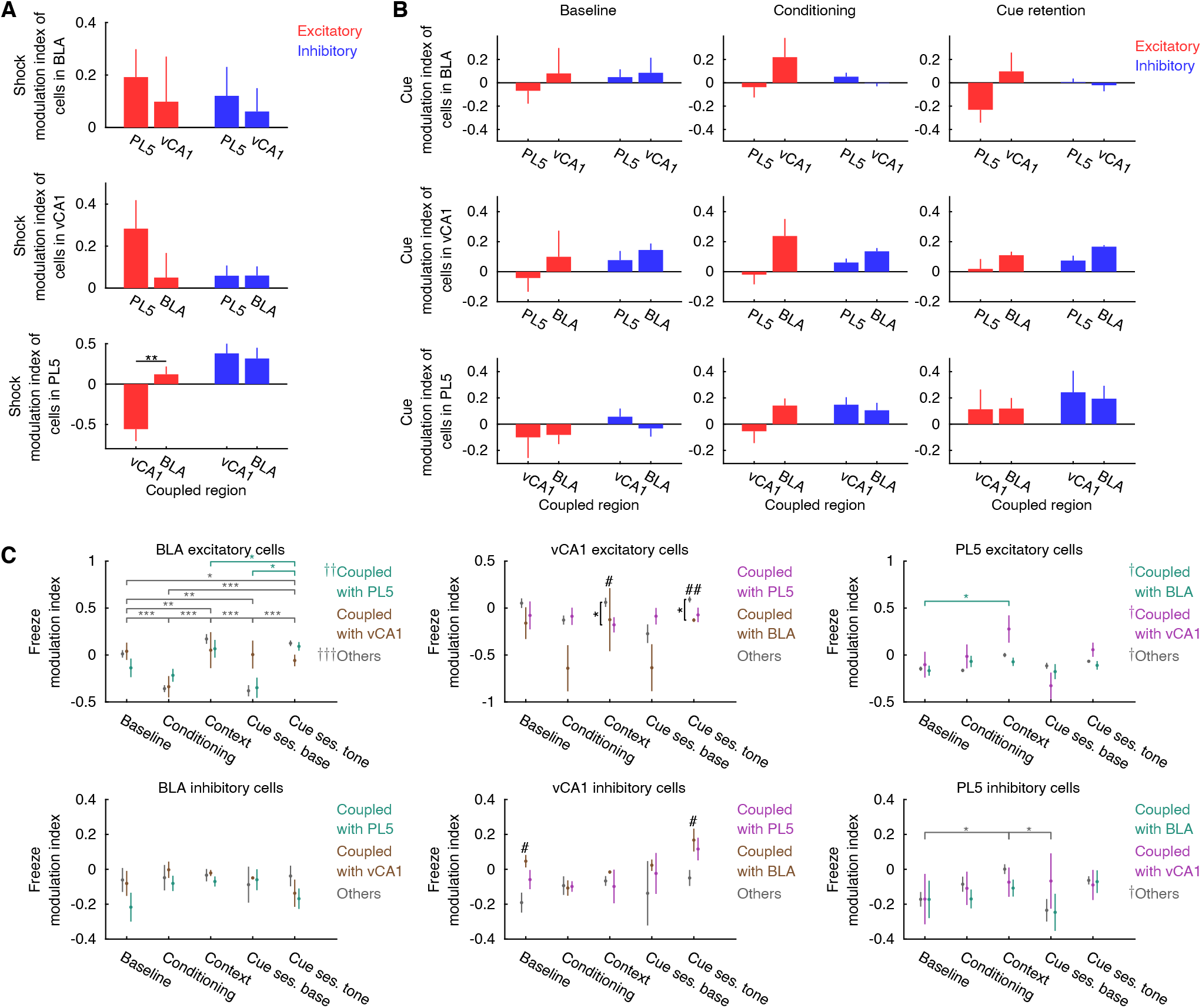
Modulation of cell firing during behavioral sessions, related to Figure 8. **(A)** Firing rate modulation of shock are compared between cells coupled with different regions in BLA, vCA1and PL5. Error bars indicate SE. ** p < 0.01, WRS-test. **(B)** Firing rate modulation of cues in baseline, conditioning, and cue retention sessions are compared between cells with inter-regional partner in vCA1, BLA, and PL5. Error bars indicate SE. **(C)** Firing rate modulation of freezing for each behavioral session. Cue retention sessions (Cue ses.) were divided into before and after the first tone onsets. Cells that were not coupled with PL5, BLA, or vCA1 are shown as others. # p < 0.05, ## p < 0.01, Kruskal–Wallis test; † p <0.05, †† p <0.01, ††† p < 0.001, Friedman test; * p < 0.05, ** p < 0.01, *** p < 0.001, post-hoc WRS-test/ WSR-test with Bonferroni correction for black/colored marks. Error bars indicate SE. Numbers of analyzed cells are summarized in Table S6.

**Table S1.**
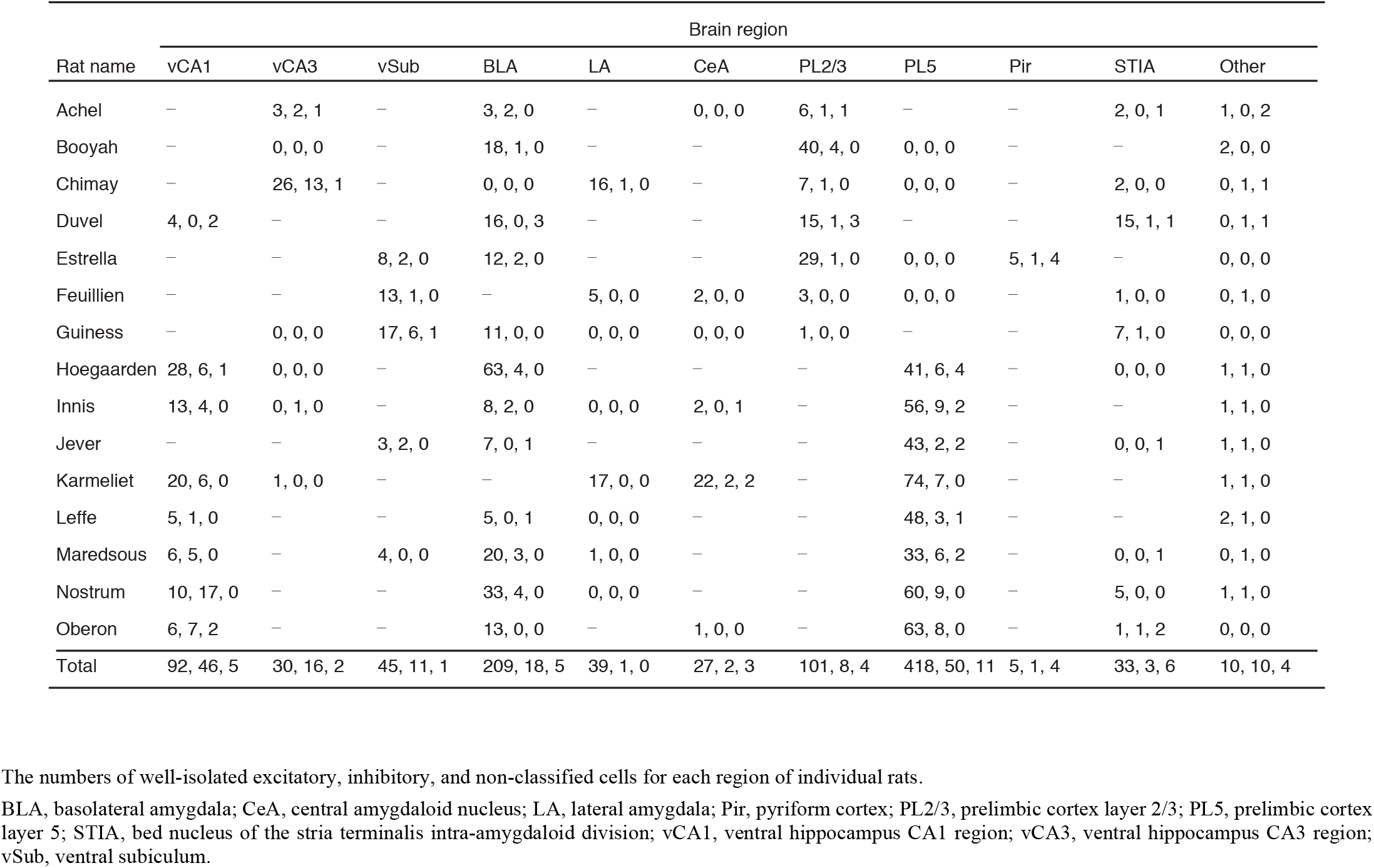
Number of recorded cells, related to Figures S1 and S3.

**Table S2.** Number of identified cell ensembles, related to Figure S3.

| Rat name | Brain region |  |  |  |  |  |  |  |  |  |  |
| --- | --- | --- | --- | --- | --- | --- | --- | --- | --- | --- | --- |
|  | vCA1 | vCA3 | vSub | BLA | LA | CeA | PL2/3 | PL5 | Pir | STIA | Other |
| Achel | – | 2 | – | 2 | – | 0 | 2 | – | – | 0 | 0 |
| Booyah | – | 0 | – | 6 | – | – | 8 | 0 | – | – | 0 |
| Chimay | – | 10 | – | 0 | 6 | – | 1 | 0 | – | 0 | 0 |
| Duvel | 1 | – | – | 3 | – | – | 5 | – | – | 4 | 0 |
| Estrella | – | – | 4 | 4 | – | – | 7 | 0 | 2 | – | 0 |
| Feuillien | – | – | 4 | – | 1 | 0 | 0 | 0 | – | 0 | 0 |
| Guinness | – | 0 | 7 | 2 | 0 | 0 | 0 | – | – | 2 | 0 |
| Hoegaarden | 8 | 0 | – | 17 | – | – | – | 10 | – | 0 | 0 |
| Innis | 5 | 0 | – | 2 | 0 | 0 | – | 12 | – | – | 0 |
| Jever | – | – | 1 | 2 | – | – | – | 8 | – | 0 | 0 |
| Karmeliet | 7 | 0 | – | – | 4 | 4 | – | 15 | – | – | 0 |
| Leffe | 2 | – | – | 2 | 0 | – | – | 8 | – | – | 0 |
| Maredsous | 3 | – | 0 | 7 | 0 | – | – | 11 | – | 0 | 0 |
| Nostrum | 7 | – | – | 9 | 0 | – | – | 14 | – | 2 | 0 |
| Oberon | 5 | – | – | 4 | – | 0 | – | 15 | – | 0 | 0 |
| Total | 38 | 12 | 16 | 60 | 11 | 4 | 23 | 93 | 2 | 8 | 0 |

The number of identified cell ensembles for each region of individual rats.
BLA, basolateral amygdala; CeA, central amygdaloid nucleus; LA, lateral amygdala; Pir, pyriform cortex; PL2/3, prelimbic cortex layer 2/3; PL5, prelimbic cortex layer 5; STIA, bed nucleus of the stria terminalis intra-amygdaloid division; vCA1, ventral hippocampus CA1 region; vCA3, ventral hippocampus CA3 region; vSub, ventral subiculum.

**Table S3.** Number of inter-regional ensemble pairs, related to Figures 3, 4, 6, S2, S4 and S6.

| Region pair | Number of pairs | Coupled pairs | Inverse-coupled pairs |
| --- | --- | --- | --- |
| BLA - PL5 | 489 | 38 (7.8%) | 10 (2.0%) |
| vCA1 - PL5 | 467 | 12 (2.6%) | 3 (0.6%) |
| vCA1 - BLA | 257 | 4 (1.6%) | 1 (0.4%) |
| BLA - Pir | 8 | 0 (0.0%) | 0 (0.0%) |
| BLA - PL2/3 | 95 | 3 (3.2%) | 2 (2.1%) |
| BLA - STIA | 34 | 1 (2.9%) | 1 (2.9%) |
| BLA - vCA3 | 4 | 0 (0.0%) | 0 (0.0%) |
| BLA - vSub | 32 | 0 (0.0%) | 0 (0.0%) |
| vCA1 - CeA | 28 | 0 (0.0%) | 0 (0.0%) |
| vCA1 - LA | 28 | 0 (0.0%) | 0 (0.0%) |
| vCA1 - PL2/3 | 5 | 0 (0.0%) | 0 (0.0%) |
| vCA1 - STIA | 18 | 0 (0.0%) | 0 (0.0%) |
| PL5 - STIA | 28 | 0 (0.0%) | 0 (0.0%) |
| PL5 - CeA | 60 | 0 (0.0%) | 0 (0.0%) |
| PL5 - LA | 60 | 0 (0.0%) | 0 (0.0%) |
| PL5 - vSub | 8 | 0 (0.0%) | 0 (0.0%) |
| vCA3 - LA | 60 | 2 (3.3%) | 3 (5.0%) |
| vCA3 - PL2/3 | 14 | 0 (0.0%) | 0 (0.0%) |
| vSub - LA | 4 | 0 (0.0%) | 0 (0.0%) |
| vSub - Pir | 8 | 0 (0.0%) | 0 (0.0%) |
| vSub - PL2/3 | 28 | 0 (0.0%) | 0 (0.0%) |
| vSub - STIA | 14 | 1 (7.1%) | 2 (14.3%) |
| LA - CeA | 16 | 0 (0.0%) | 0 (0.0%) |
| LA - PL2/3 | 6 | 0 (0.0%) | 0 (0.0%) |
| PL2/3 - Pir | 14 | 0 (0.0%) | 0 (0.0%) |
| PL2/3 - STIA | 20 | 1 (5.0%) | 1 (5.0%) |

The number of simultaneously recorded inter-regional ensemble pairs and number and fraction of coupled and inverse-coupled ensemble pairs. Chance levels for the fractions of coupled and inverse-coupled ensemble pairs were 0.5%.
BLA, basolateral amygdala; CeA, central amygdaloid nucleus; LA, lateral amygdala; Pir, pyriform cortex; PL2/3, prelimbic cortex layer 2/3; PL5, prelimbic cortex layer 5; STIA, bed nucleus of the stria terminalis intra-amygdaloid division; vCA1, ventral hippocampus CA1 region; vCA3, ventral hippocampus CA3 region; vSub, ventral subiculum.

**Table S4.**
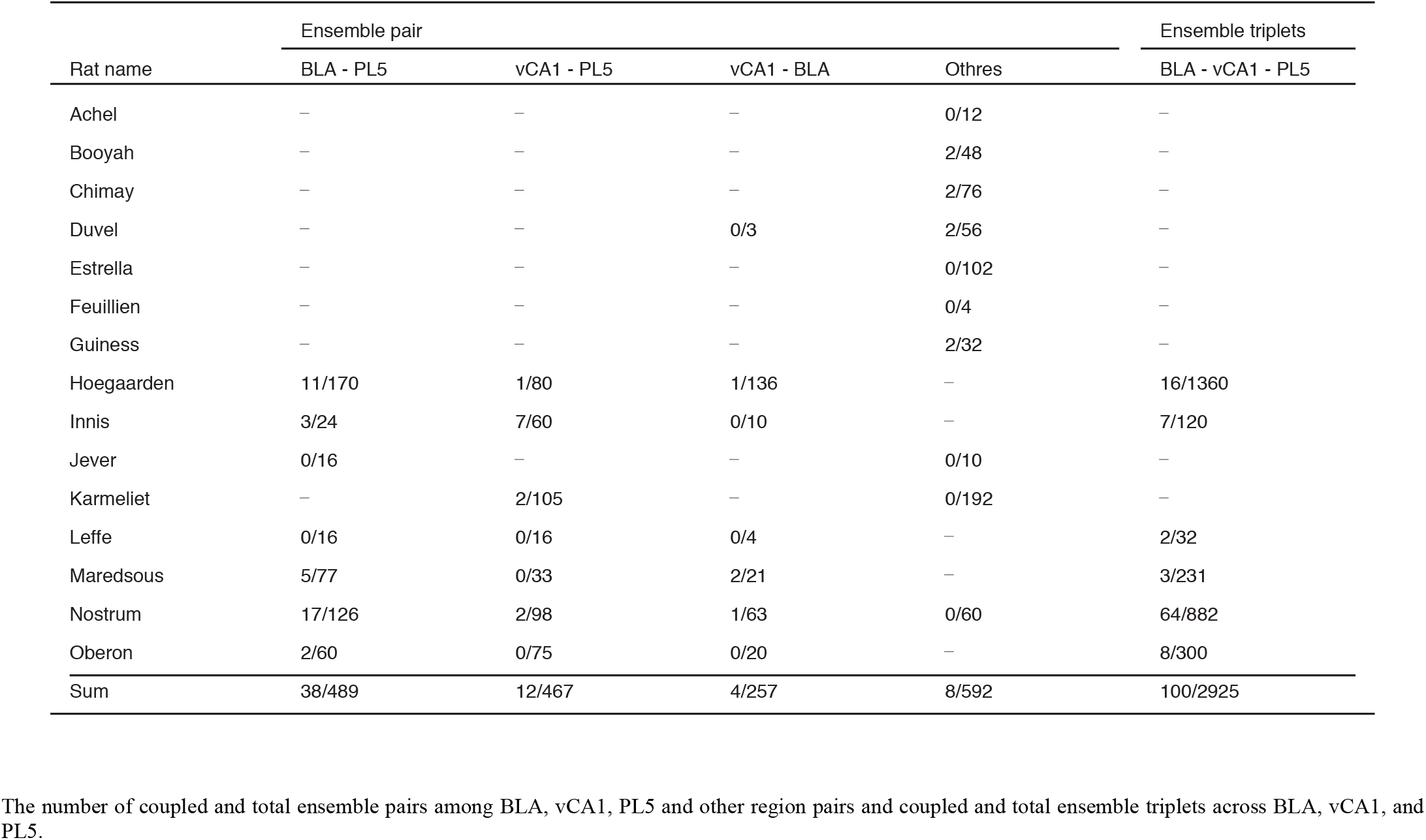
Number of coupled ensemble pairs and triplets in each rat, related to Figures 3 and 5.

**Table S5.** Number of ensembles, related to Figures 8.

|  | Coupled with BLA | Coupled with vCA1 | Coupled with PL5 | Other ensembles |
| --- | --- | --- | --- | --- |
| Ensembles in BLA | N.A. | 4 | 15 | 44 |
| Ensembles in vCA1 | 3 | N.A. | 8 | 28 |
| Ensembles in PL5 | 22 | 7 | N.A. | 68 |

The number of ensembles coupled with ensembles in other brain regions. Ensembles that were not coupled with BLA, vCA1 or PL5 are shown as other ensembles. NA, not applicable.

**Table S6.** Number of cells highly contributing to neuronal ensembles, related to Figures 8, S7, and S8.

|  | Coupled with BLA | Coupled with vCA1 | Coupled with PL5 | Other cells |
| --- | --- | --- | --- | --- |
| Cells in BLA | N.A. | 10 / 5 / 0 | 36 / 10 / 0 | 169 / 8 / 5 |
| Cells in vCA1 | 5 / 5 / 0 | N.A. | 13 / 13 / 0 | 74 / 31 / 5 |
| Cells in PL5 | 37 / 12 / 2 | 10 / 6 / 0 | N.A. | 373 / 38 / 9 |

The number of excitatory/inhibitory/non-classified cells that highly contributed to ensembles coupled with other brain regions. Cells that were not coupled with BLA, vCA1 or PL5 are shown as other cells. NA, not applicable.

